# Dynamics of Cardiomyocyte Transcriptome and Chromatin Landscape Demarcates Key Events of Heart Development

**DOI:** 10.1101/488593

**Authors:** Michal Pawlak, Katarzyna Z. Kedzierska, Maciej Migdal, Karim Abu Nahia, Jordan A. Ramilowski, Lukasz Bugajski, Kosuke Hashimoto, Aleksandra Marconi, Katarzyna Piwocka, Piero Carninci, Cecilia L. Winata

## Abstract

The development of an organ involves dynamic regulation of gene transcription and complex multipathway interactions. To better understand transcriptional regulatory mechanism driving heart development and the consequences of its disruption, we isolated cardiomyocytes (CMs) from wild-type zebrafish embryos at 24, 48 and 72 hours post fertilization corresponding to heart looping, chamber formation and heart maturation, and from mutant lines carrying loss-of-function mutations in *gata5, tbx5a* and *hand2*, transcription factors (TFs) required for proper heart development. The integration of CM transcriptomics (RNA-seq) and genome-wide chromatin accessibility maps (ATAC-seq) unravelled dynamic regulatory networks driving crucial events of heart development. These networks contained key cardiac TFs including Gata5/6, Nkx2.5, Tbx5/20, and Hand2, and are associated with open chromatin regions enriched for DNA sequence motifs belonging to the family of the corresponding TFs. These networks were disrupted in cardiac TF mutants, indicating their importance in proper heart development. The most prominent gene expression changes, which correlated with chromatin accessibility modifications within their proximal promoter regions, occurred between heart looping and chamber formation, and were associated with metabolic and hematopoietic/cardiac switch during CM maturation. Furthermore, loss of function of cardiac TFs Gata5, Tbx5a, and Hand2 affected the cardiac regulatory networks and caused global changes in chromatin accessibility profile. Among regions with differential chromatin accessibility in mutants were highly conserved non-coding elements which represent putative *cis* regulatory elements with potential role in heart development and disease. Altogether, our results revealed the dynamic regulatory landscape at key stages of heart development and identified molecular drivers of heart morphogenesis.

## INTRODUCTION

The heart muscle or myocardium makes up most of the heart tissues and is mainly responsible for its function. Upon completion of gastrulation, heart muscle cells or cardiomyocytes (CMs) are specified from a pool of mesodermal progenitors at the anterior portion of the embryonic lateral plate mesoderm (Stainier et al. 1993; Stainier and Fishman 1994; Kelly et al. 2014). As development proceeds, heart progenitors migrate to the midline and form a tube structure known as the primitive heart tube (Stainier et al. 1993). This structure subsequently expands through cell division and addition of more cells originating from the progenitor pool (Kelly et al. 2014; Knight and Yelon 2016). Looping of the heart tube then gives rise to distinct chambers of the heart, namely, the atria and ventricles. Although the vertebrate heart can have between two to four chambers, the step-wise morphogenesis of progenitors specification, migration, tube formation, and looping, are highly conserved between species (Jensen et al. 2013).

CMs are specified early during embryogenesis and undergo various cellular processes of proliferation, migration, and differentiation which collectively give rise to a fully formed and functioning heart. Crucial to regulating each step of heart morphogenesis are cardiac transcription factors (TFs) which include Nkx2.5, Gata5, Tbx5, and Hand2 (Clark et al. 2006; Nemer 2008). These TFs are known to play a role in establishing the CM identity of mesodermal progenitor cells, regulating the formation and looping of the heart tube, as well as the specification of atrial and ventricular CMs. Specification and differentiation of the cardiac progenitors are regulated by the interactions between several key TFs - Nkx2.5, Gata5, Tbx5, and Hand2. Members of the GATA family of TFs, Gata4, Gata5, and Gata6, are responsible for the earliest step of cardiac progenitor specification (Jiang and Evans 1996; Jiang et al. 1998; Reiter et al. 1999; Singh et al. 2010; Lou et al. 2011; Turbendian et al. 2013). Gata factors activate the expression of *nkx2.5*, another early marker of CMs (Chen and Fishman 1996; Lien et al. 1999). Although not essential for specification of CMs, Nkx2.5 plays an important role in initiating the expression of many cardiac genes in mouse and regulating the numbers of atrial and ventricular progenitors (Searcy et al. 1998; Targoff et al. 2008). Similarly, another TF expressed in CM progenitors, Hand2, is responsible for proliferation of ventricular progenitors (Yelon et al. 2000). Hand2 also induces and maintains the expression of Tbx5, which is necessary for atrial specification in the mouse (Liberatore et al. 2000; Bruneau et al. 2001).

Despite the established knowledge of key TFs regulating the various steps of heart morphogenesis, considerable challenges to understand the mechanism of heart development still exist as little is known about their molecular mechanism and downstream targets. Transcription is modulated by *cis* regulatory elements that are located in non-coding regions of the genome, which serve as binding sites for TFs (Farnham 2009; Shlyueva et al. 2014). Although these regulatory elements equally contribute to the molecular mechanism controlling development, there is still a lack of systematic resources and understanding of their roles in heart development. Moreover, cardiac TFs have been shown to interact with chromatin-modifying factors, and the loss of function of several histone-modifying enzymes has been found to affect various aspects of cardiac development (Miller et al. 2008; Nimura et al. 2009; Lou et al. 2011; Takeuchi et al. 2011). Therefore, the chromatin landscape is another factor which needs to be considered when studying the process of heart development. Importantly, the lack of understanding how heart development proceeds makes it difficult to determine the cause of different forms of congenital heart disease (CHD). Here we seek to understand the nature of interaction between TFs and epigenomic landscape, how this landscape changes throughout development, and how it affects heart development.

The study of heart development poses a unique challenge due to the importance of the organ for survival. The disruption of factors regulating the early steps of heart formation can result in early embryonic lethality. The use of zebrafish as a model organism alleviates this problem by allowing access to developing embryos immediately after fertilization and its ability to survive without a functioning heart up to a comparatively late stage of development (Stainier 2001; Staudt and Stainier 2012). To elucidate the dynamics of the transcriptional regulatory landscape during heart development, we isolated CMs directly from the developing wild-type zebrafish heart at three key stages of morphogenesis: linear heart tube formation (24 hpf), chamber formation and differentiation (48 hpf), and heart maturation (72 hpf). Similarly, we isolated CMs from cardiac TF mutants of *gata5, tbx5a* and *hand2* at 72 hpf. We then combined transcriptome profiling (RNA-seq) with an assay for chromatin accessibility (ATAC-seq) (Buenrostro et al. 2013) to capture the dynamics of regulatory landscape throughout the progression of heart morphogenesis *in vivo*. Our results unravelled the gene regulatory network driving key processes of heart development.

## RESULTS

### CM transcriptome reveals strong dynamics at early stages of heart morphogenesis

One of the earliest markers of cardiac lineage are NK2 homeobox 5 (*nkx2.5*), which is expressed in cardiac precursor cells in the anterior lateral plate mesoderm and is required in the second heart field as the heart tube forms (George et al. 2015), and myosin light chain 7 (*myl7*), responsible for sarcomere assembly and specific to differentiated myocardial cells (Chen et al. 2008). To study gene regulatory networks underlying zebrafish heart development, we isolated CMs from zebrafish transgenic lines Tg(*nxk2.5*:GFP) (Witzel et al. 2012) and Tg(*myl7*:EGFP) (D’Amico et al. 2007) using fluorescence-activated cell sorting (FACS, Fig. 1A). Cells were collected at three different stages of heart development which corresponded to linear heart tube formation (24 hpf), chamber formation and differentiation (48 hpf) and adult heart maturation (72 hpf) (Bakkers 2011) (Fig. 1B). Due to its earlier onset of CM-specific GFP expression Tg(*nxk2.5*:GFP) were used to sort CM at 24 hpf, whereas Tg(*myl7*:EGFP) were used for the subsequent developmental stages (48 hpf and 72 hpf)(Houk and Yelon 2016). The average fraction of FACS-yielded GFP+ events obtained from embryo cell suspension at all three stages of development were between 1.37 to 2.56% of total singlet events (Supplement. Fig. 1A). To monitor the purity of FACS and establish the identity of the isolated cells, we measured mRNA levels of *nkx2.5, myl7* and GFP in both GFP+ and GFP-cells. The expression of the CM markers and GFP were strongly enriched in GFP+ as compared to GFP-fraction (Supplement. Fig. 1B). In contrast, mRNA levels of *neurogenini* (*ngnl*), a neuronal-specific gene, were higher in GFP-cells. In line with that, RNA-seq expression of *nkx2.5, myl7* and *myh6* was strongly enriched in GFP+ as compared to GFP-cells, whereas expression of non-CM markers such as skeletal muscle (*myog*), pancreas (*ins*), pharyngeal arch (*frem2a*), retina (*arr3b, otx5*), skin (*tp63, col16a1*), neural system (*neurog1, zic3, otx1*) and eye (*pou4f2*) was more pronounced in GFP-(Supplement. Fig. 2). Additionally, RNA-seq followed by gene ontology (GO) enrichment analysis of differentially expressed genes between GFP+ and GFP-across all three stages of heart development revealed the overrepresentation of CM-specific biological processes such as cell migration, cardiac development and heart function (Fig. 1C, Supplement Table 1). Among 50 genes with the highest average expression across all developmental stages, 35 are known to have specific functions in CM according to ZFIN database (https://zfin.org) and eight are associated with CM-specific functions and human diseases such as cardiac muscle contraction and cardiomyopathy (*ttn.1, mybpc3, ttn.2, acta1b, actn2b*), atrial septal defects (*actc1a, myh6*) and Laing distal myopathy (*vmhc*) (Fig. 1D) according to the Online Mendelian Inheritance in Man (OMIM) database (https://www.omim.org/).

**Figure 1.**
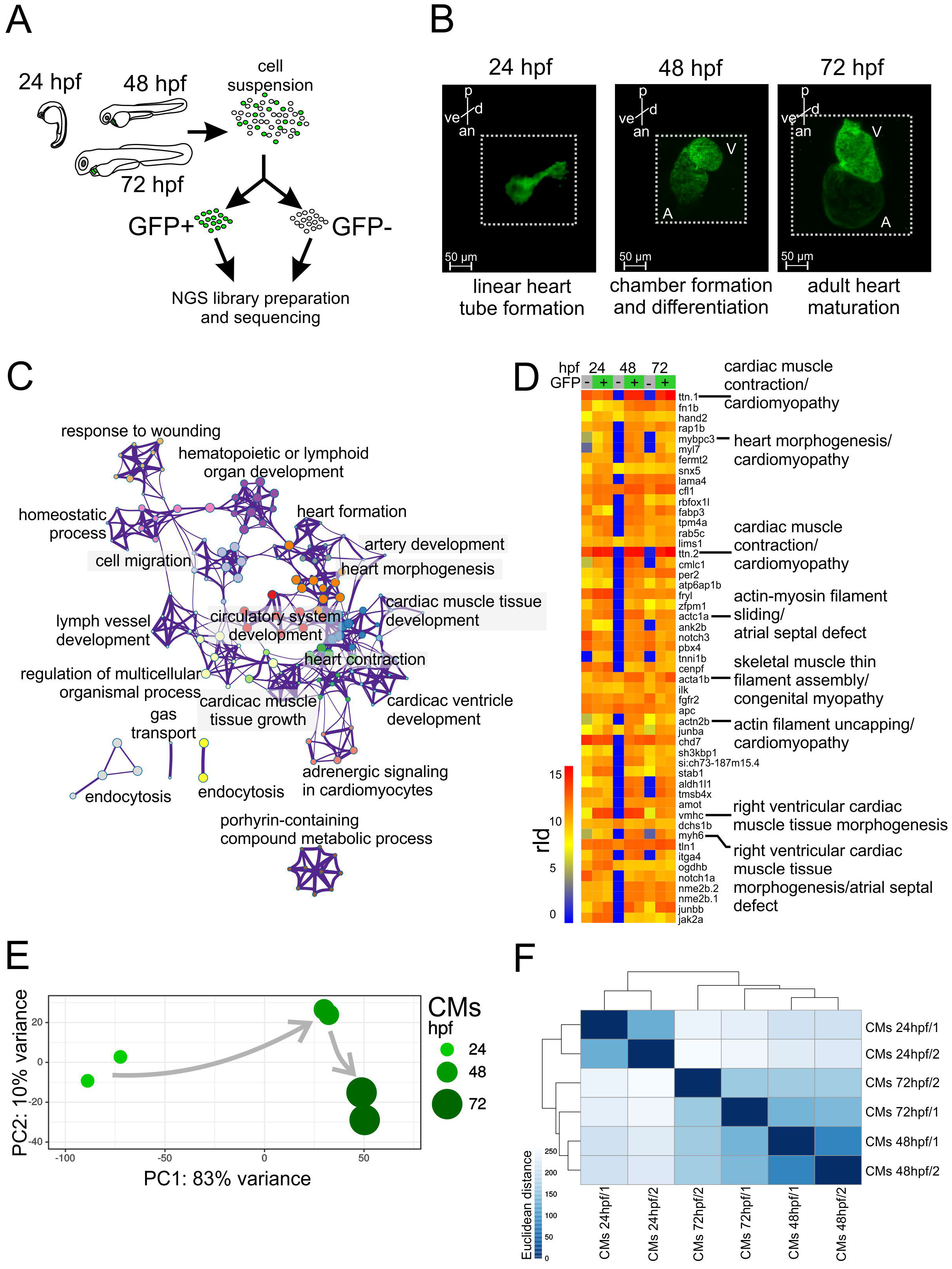
CM transcriptome landscape during heart development. **(A)** Schematics of experimental design. **(B)** Light sheet fluorescence microscope (LSFM) images of GFP-labeled CMs of developing zebrafish heart. p - posterior, an - anterior, v - ventral, d – dorsal. Dotted line indicates exact area of the LSFM image. **(C)** Network of 20 top-score GO clusters enriched in genes commonly upregulated in GFP+ across heart development. Size nodes refer to the number of genes contributing to the same GO and nodes that share the same cluster ID are close to each other, padj ≤ 0.05. **(D)** Heatmap of top 50 highly expressed genes between 24-72 hpf based on normalized expression value (regularized log, rld). **(E)** Graphical representation of PCA of CM RNA-seq data. **(F)** Heatmap and clustering of RNA-seq sample-to-sample Euclidean distances.

To explore the dynamics of zebrafish CM transcriptome during heart development we applied principal component analysis (PCA) and RNA-seq sample clustering based on Euclidean distance (see Methods). Both analyses revealed strong dissimilarity in transcriptome profiles between CM at 24 hpf and later stages of heart development. This suggest that the major gene expression profile changes occur in CM between 24 and 48 hpf and correspond to linear heart tube formation and chamber formation as compared to CM at 48 and 72 hpf which showed stronger similarity (Fig. 1E-F).

Taken together, we have successfully isolated CMs from zebrafish heart *in vivo* at three developmental stages. Our transcriptome analyses identified CM-specific gene expression signatures among highly abundant transcripts and revealed the dynamic nature of gene expression profiles during the course of heart morphogenesis.

### Chromatin accessibility is correlated with CM gene expression levels during heart development

The chromatin landscape, in combination with TF-mediated regulation, is known to control cell differentiation and organ development (He et al. 2014; Karwacz et al. 2017; Nelson et al. 2017). To characterize chromatin dynamics throughout heart development, we used assay for transposase accessible chromatin with high-throughput sequencing (ATAC-seq) and profiled chromatin accessibility at three developmental stages matching our transcriptome analyses: 24 hpf, 48 hpf, and 72 hpf (Buenrostro et al. 2013). To identify genome-wide nucleosome free regions (NFR), ATAC-seq read fragments were partitioned into four populations (Fig. 2A) based on exponential function for fragment distribution pattern at insert sizes below one nucleosome (123 bp) and Gaussian distributions for 1, 2 and 3 nucleosomes as previously described (Buenrostro et al. 2013). The PCA analysis of (Fig. 2B) and clustering using the Euclidian distances between ATAC-seq samples based on their NFR profiles (Fig. 2C) revealed that biological replicas clustered together, whereas, the largest changes in chromatin accessibility were observed between 24 hpf and 48 hpf stages, in agreement with observed transcriptome changes of CMs during heart development. Comparing consensus NFRs across all developmental stages, we observed a large number of common NFRs (16,055), as well as those which were specific to a single developmental stage. The most stage-specific NFRs were found in CMs at 24 hpf (22,656) (Fig. 2D). This prompted us to further investigate the relationship between transcriptome and chromatin accessibility changes in cardiac development. We therefore looked at the distribution of NFRs across genomic features and observed that the highest fraction of NFRs was localized either within promoter regions (∼30% of total NFRs), followed by intergenic (∼25%) and intronic (20%) regions (Fig. 2E, Supplement Table 2). These ratios remained at comparable levels across all three developmental stages studied. Consistently, NFR consensus heatmaps within transcription start site (TSS) proximal promoter regions (+/- 3 kb) (Fig. 2F) compared to distal promoter regions (more than +/- 3 kb of TSS) (Fig. 2G) as well as ATAC-seq read density over the gene bodies of 1000 genes most highly expressed in CMs at all three stages of heart development (Fig. 2H) revealed the enrichment of NFRs around TSS regions. We further observed that chromatin accessibility reflected by the presence of NFR in gene promoter regions was significantly correlated with the expression levels of the corresponding genes to which the promoter belonged to (Spearman rho 0.46 - 0.48) at each stage of heart development (Fig. 2I). Our observations therefore revealed a strong link between chromatin accessibility of promoter regions and gene expression levels.

**Figure 2.**
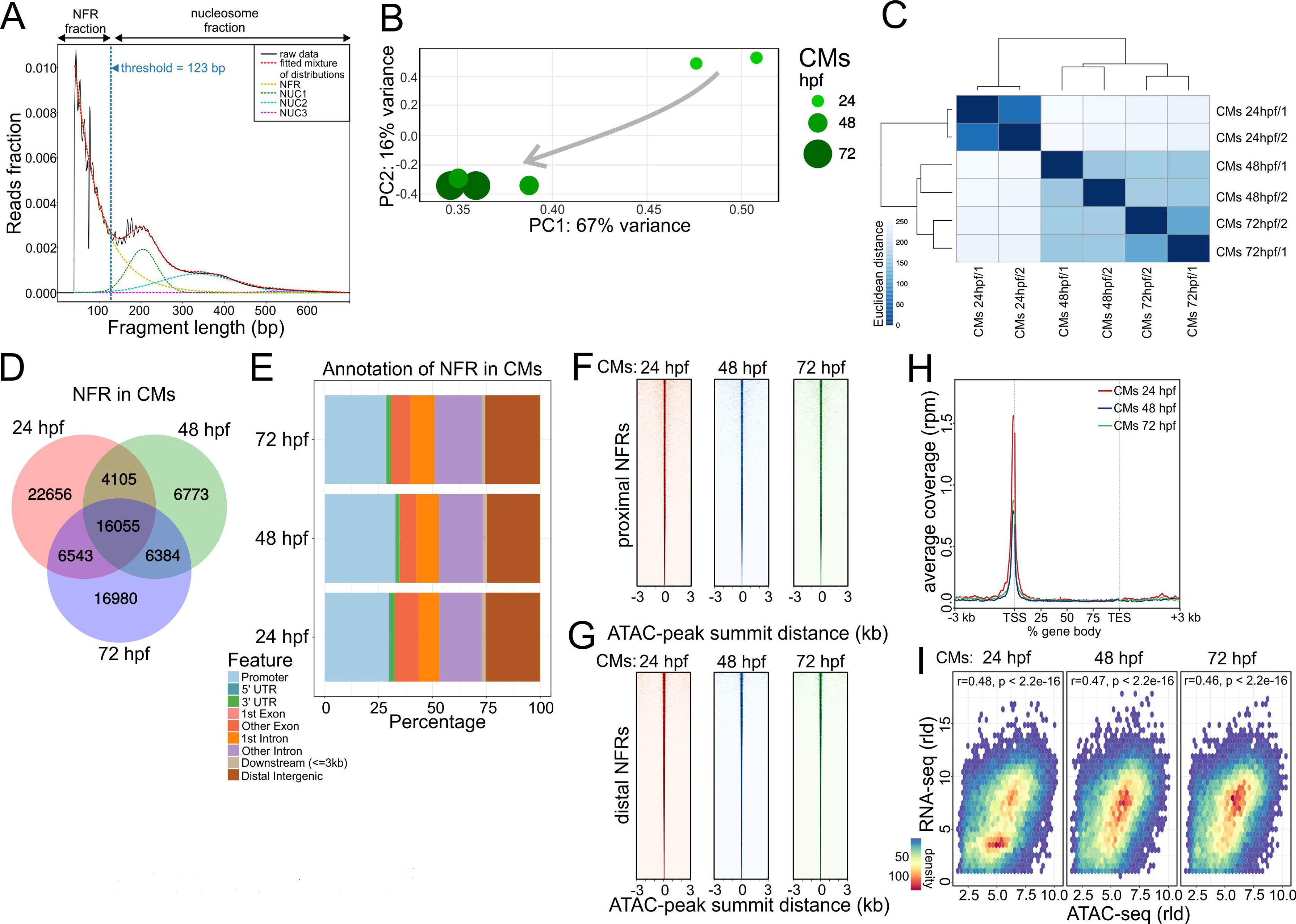
Cross-talk between transcriptome and chromatin accessibility profile across stages of cardiac development. **(A)** ATAC-seq read distribution and characterization of NFR fractions. **(B)** PCA of NFR chromatin accessibility during heart development. **(C)** Euclidian distances between chromatin accessibility within NFR. **(D)** Comparison of NFR presence and overlap across stages of heart development. **(E)** Genomic annotation of CM NFR consensus at different stages of heart development. **(F)** CM NFR consensus coverage heatmap of TSS proximal (+/-3kb of TSS) regions centred on ATAC-seq peak summits. (**G)** CM NFR consensus coverage heatmap of TSS distal (more than +/-3kb of TSS) regions centred on ATAC-seq peak summits. **(H)** Metaplot of ATAC-seq read density over the gene bodies of 1000 genes most highly expressed in CMs at each developmental stage. TES – transcription end site. **(I)** Spearman correlation of normalized log (rld) RNA-seq gene expression and ATAC-seq chromatin accessibility in corresponding NFR regions (+/-3kb of TSS).

### Co-expression network analysis identifies CM regulatory modules

Analysis of transcriptional profiles across different conditions allows to organize genes with similar expression patterns into functional regulatory modules (Langfelder and Horvath 2008). To better understand the relationship and functionality of cardiac genes involved in the developing zebrafish heart *in vivo*, we identified relevant gene regulatory networks in an unsupervised and unbiased manner using the weighted gene correlation network analysis (WGCNA) based on RNA-seq expression profiles (Langfelder and Horvath 2008). Hierarchical clustering of the similarity/dissimilarity matrix across the entire set of transcriptome samples distinguished 37 gene modules (Fig. 3A, Supplement Table 3), out of which five were enriched in functional terms related to cardiovascular system development and function (Fig. 3B, Supplement Table 4): turquoise (4085 genes), brown (2156 genes), green (1166 genes), salmon (756 genes), and sienna3 (75 genes). We refer to these modules as “cardiac modules” from here on. Functional terms enriched in these cardiac modules included specific processes of heart development, such as “embryonic heart tube development” (modules brown, green, and sienna3), “cardioblast differentiation” (green), “heart valve development” (salmon), “heart process” and “heart formation” (turquoise). The relatively small sienna3 module was strongly enriched in GO terms associated with multiple cardiac developmental processes including “heart tube development”, “cardioblast migration” and “heart rudiment development”.

**Figure 3.**
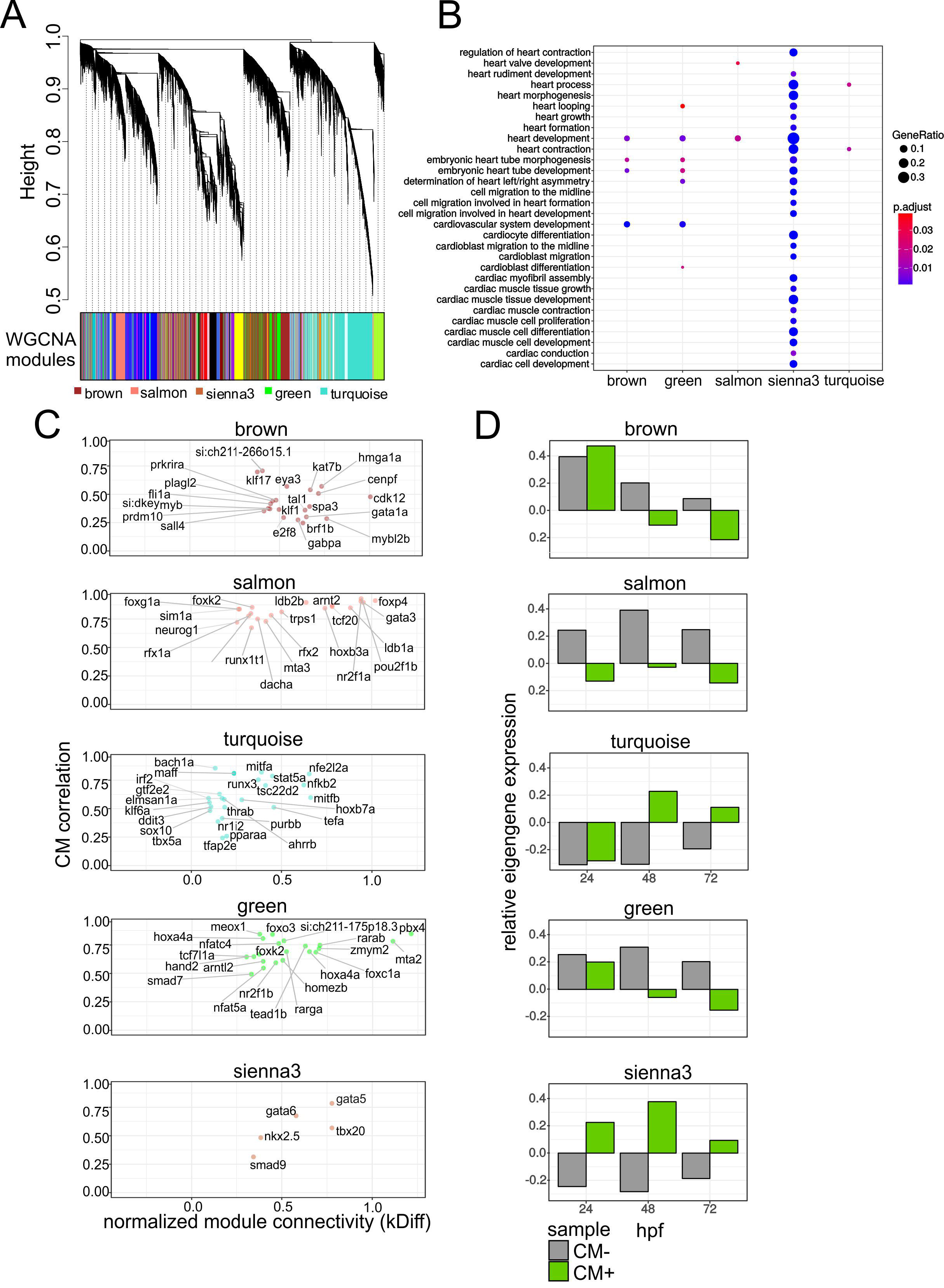
Cardiac co-expression regulatory networks. **(A)** Hierarchical clustering of gene expression similarity/dissimilarity matrix. **(B)** Cardiovascular-related GO enrichment in five cardiac modules. **(C)** Module gene connectivity plot of selected TFs. Twenty TFs with the highest normalized kDiff are shown. **(D)** Cardiac module eigengene expression during heart development.

To unravel potential driver genes with regulatory roles in each of the cardiac modules identified, we searched for transcription factors (TFs) and calculated their connectivity to other genes within a given module (normalized kDiff), as well as how their expression is affected by a CM phenotypic trait (CM correlation) (Fig. 3C). Most of the cardiac modules contained TFs known to direct key processes of heart development, such as *gata1* (brown), *tbx5a, sox10* (turquoise), *hand2, smad7* (green) as well as *gata5, nkx2.5, tbx20* (sienna3) (Reiter et al. 1999; Ahn et al. 2000; Montero et al. 2002; Holtzinger and Evans 2007; Schoenebeck et al. 2007; Targoff et al. 2008; Moskowitz et al. 2011; Ounzain et al. 2014). Each of the modules exhibited different expression profile dynamics in heart development (termed eigengene expression) across three developmental stages in both GFP+ and GFP-fraction, further called CM+ and CM-, respectively (Fig 3D). Two broad patterns of eigengene expression could be observed: modules with decreasing cardiac gene expression during heart development - brown and green, and modules in which expression increases between 24 and 48 hpf and then decreases between 48 and 72 hpf - salmon, sienna3 and turquoise. In addition, CM+ eigengene expression in sienna3 module was consistently higher than in CM-samples at all stages of development, further suggesting the specificity of this module to CM.

The presence of key cardiac TFs in each module prompted us to look closer into individual genes within these modules so to identify specific functional patterns related to cardiovascular development. The sienna3 module, which contained cardiac TFs *nkx2.5, gata5, gata6*, and *tbx20*, also contained many other genes implicated in various aspects of heart morphogenesis including CM migration and differentiation, and heart looping including *popdc2, apobec2a*, and *tdgf1* (Xu et al. 1999; Kirk et al. 2007; Etard et al. 2010; Wang et al. 2011; Kirchmaier et al. 2012; Sakabe et al. 2012). Additionally, the module also contained many genes known to be involved in cell adhesion and structural constituents of the heart muscle, which were previously implicated in cardiomyopathy when mutated. These included *actc1a, myl7, myh7ba, myh7bb, vmhc, and ttn.2* (Olson et al. 1998; Xu et al. 2002; Shih et al. 2015). In support of this network, *popdc2* and *gata6* were previously shown to be a direct transcriptional target of Nkx2.5 in mouse embryonic heart (Davis et al. 2000; Molkentin et al. 2000; Dupays et al. 2015). In turn, evidence also exists for the cardiac-specific transcriptional activation of *nkx2.5* by GATA factors (Lien et al. 1999).

Genes belonging to the developmental signaling pathways Wnt, Notch, TGF-◻ and FGF were highly represented in all modules except sienna3 which consisted of mostly specialized CM genes. In particular, genes of both canonical and non-canonical Wnt signaling pathways were almost exclusively distributed between the green and salmon modules. Studies in different organism have shown that the canonical Wnt signaling plays biphasic roles in cardiac development, where it promotes cardiac fate in the early precardiac mesoderm while becoming inhibitory to cardiogenesis processes in later stages (Naito et al. 2006; Ueno et al. 2007; Piven and Winata 2017). The cardiac TF Hand2, known to regulate early cardiac developmental processes, is also present in the green module, suggesting that it might control these pathways. Altogether, we identified regulatory modules exhibiting unique expression patterns throughout heart development, each of which contained relevant TFs. Importantly, these modules represent potential regulatory networks underlying various processes of heart development.

### Integrative analysis of RNA-seq and ATAC-seq identifies regulatory networks of CM maturation

To further explore the relationship between chromatin state and transcriptional regulation of heart development, we integrated co-expression networks generated from RNA-seq with accessible chromatin regions identified by ATAC-seq. Thus, we examined NFRs localized within +/-3kb of the TSS of genes assigned to the same module for the presence of TF motifs (Table 1, Supplement Fig. 3). NFRs associated with genes within the module sienna3 (which contained Gata5/6, Nkx2.5, and Tbx20 TFs) were also enriched in motifs belonging to these family of TFs [Gata family (Gata1/2/3/4/6), Nkx family (Nkx2.2, Nkx2.5), Smad3 and T-box family (Tbr1)], whereas salmon module containing *sox3* gene showed overrepresentation of Sox3 motif. Similarly, in two other cardiac modules turquoise and green (containing the TFs Tbx5, Hand2, and Smad7) we found a wide range of significantly enriched (pvalue ≤ 0.05) TF motifs including Tbx family (Tbx5) and Smad family (Smad2, Smad4), respectively. The presence of the TFs together with the enrichment of their respective recognition motifs strongly suggests their regulatory role within each module. Moreover, we observed an overrepresentation of motifs of TF with profound role in heart development, such as Sox family (Sox10) motifs in salmon module and Tgif family (Tgif1, Tgif2) in both sienna3 and turquoise modules (Montero et al. 2002; Powers et al. 2010) although TFs corresponding to these motifs were not present in the matching modules.

**Table 1.** HOMER-identified TF motifs found in NFR of cardiac co-expression modules. HOMER-identified motifs with the highest prevalence in NFRs localized +/-3kb around the TSSs of selected cardiac module genes are listed. P-value < 0.05. Known vertebrate TF motifs were used for analysis.

| brown |  |  |  |  | green |  |  |  |  |
| --- | --- | --- | --- | --- | --- | --- | --- | --- | --- |
| Motif Name | P-value | Target Sequences with Motif (of 4698) | % of Target Sequences with Motif | % of Background Sequences with Motif | Motif Name | P-value | Target Sequences with Motif (of 4698) | % of Target Sequences with Motif | % of Background Sequences with Motif |
| Smad3 | 1.E-02 | 1172 | 24.94% | 23.07% | Smad4 | 1.E-04 | 545 | 19.93% | 16.96 |
| bHLH | 1.E-02 | 904 | 19.24% | 17.83% | Smad2 | 1.E-02 | 495 | 18.11% | 15.85 |
| Fli1 | 1.E-09 | 786 | 16.73% | 13.44% | Sox3 | 1.E-02 | 420 | 15.36% | 13.41 |
| ETV1 | 1.E-05 | 729 | 15.51% | 12.97% | Sox6 | 1.E-02 | 407 | 14.89% | 12.87 |
| NFY | 1.E-06 | 702 | 14.94% | 12.41% | Sox10 | 1.E-02 | 387 | 14.16% | 12.41 |
| Sox3 | 1.E-02 | 700 | 14.90% | 13.63% | Gata4 | 1.E-02 | 284 | 10.39% | 8.84 |
| ERG | 1.E-03 | 672 | 14.30% | 12.46% | Gata6 | 1.E-02 | 255 | 9.33% | 7.79 |
| GATA3 | 1.E-03 | 664 | 14.13% | 12.30% | Sox2 | 1.E-02 | 218 | 7.97% | 6.7 |
| ETS1 | 1.E-07 | 619 | 13.17% | 10.60% | Sox4 | 1.E-03 | 212 | 7.75% | 6.12 |
| EHF | 1.E-02 | 581 | 12.36% | 11.16% | Gata2 | 1.E-03 | 201 | 7.35% | 5.75 |
| salmon |  |  |  |  | sienna3 |  |  |  |  |
| Motif Name | P-value | Target Sequences with Motif (of 4698) | % of Target Sequences with Motif | % of Background Sequences with Motif | Motif Name | P-value | Target Sequences with Motif (of 4698) | % of Target Sequences with Motif | % of Background Sequences with Motif |
| Sox3 | 1.E-08 | 332 | 18.24% | 13.21% | Tgif1 | 1.E-02 | 77 | 45.03% | 33.41% |
| Sox10 | 1.E-07 | 307 | 16.87% | 12.34% | Tgif2 | 1.E-02 | 77 | 45.03% | 34.57% |
| NeuroG2 | 1.E-02 | 299 | 16.43% | 14.17% | Meis1 | 1.E-02 | 46 | 26.90% | 18.63% |
| Sox6 | 1.E-04 | 293 | 16.10% | 12.60% | Nkx2.5 | 1.E-02 | 44 | 25.73% | 17.61% |
| Atoh1 | 1.E-02 | 215 | 11.81% | 9.78% | Bapx1 | 1.E-02 | 42 | 24.56% | 16.10% |
| Sox15 | 1.E-06 | 202 | 11.10% | 7.65% | Nkx2.2 | 1.E-02 | 41 | 23.98% | 16.70% |
| Sox2 | 1.E-06 | 178 | 9.78% | 6.64% | GATA3 | 1.E-03 | 38 | 22.22% | 13.17% |
| NeuroD1 | 1.E-02 | 159 | 8.74% | 7.17% | Mef2b | 1.E-09 | 33 | 19.30% | 5.66% |
| Sox4 | 1.E-04 | 157 | 8.63% | 6.08% | Tbr1 | 1.E-02 | 33 | 19.30% | 12.02% |
| Maz | 1.E-02 | 141 | 7.75% | 6.03% | Gata6 | 1.E-03 | 29 | 16.96% | 8.61% |
| turquoise |  |  |  |  |  |  |  |  |  |
| Motif Name | P-value | Target Sequences with Motif (of 4698) | % of Target Sequences with Motif | % of Background Sequences with Motif |  |  |  |  |  |
| SCL | 1.E-06 | 3906 | 45.22% | 42.29% |  |  |  |  |  |
| Tgif2 | 1.E-20 | 3467 | 40.14% | 34.68% |  |  |  |  |  |
| Tgif1 | 1.E-20 | 3353 | 38.82% | 33.42% |  |  |  |  |  |
| Nanog | 1.E-04 | 3311 | 38.34% | 35.88% |  |  |  |  |  |
| Pitx1 | 1.E-08 | 3055 | 35.37% | 31.99% |  |  |  |  |  |
| THRb | 1.E-03 | 2376 | 27.51% | 25.78% |  |  |  |  |  |
| Tbx5 | 1.E-06 | 2270 | 26.28% | 23.71% |  |  |  |  |  |
| Nkx6.1 | 1.E-02 | 2166 | 25.08% | 23.66% |  |  |  |  |  |
| AR-halfsite | 1.E-02 | 2120 | 24.55% | 23.18% |  |  |  |  |  |
| Smad3 | 1.E-02 | 2076 | 24.04% | 22.37% |  |  |  |  |  |

To establish the relationship between chromatin accessibility and gene expression and provide the link between TF and their effector genes, we combined gene-to-gene correlation with NFR motif annotation and its accessibility within the proximity (+/- 3 kb) of their transcription start site (TSS) (Fig. 4A). To identify genes which were dynamically regulated and associated with regions with differential chromatin accessibility in the course of heart development, we compared normalized changes of gene expression to those of the corresponding NFRs between 24 and 48 hpf as well as 48 and 72 hpf (Fig. 4B, Supplement Table 1 and 5). We observed strong up-regulation of expression for a large number of genes within the turquoise and salmon module and down-regulation of genes in brown module and for most genes belonging to the green module. This was generally consistent with the direction of changes in chromatin accessibility e.g. *gpd2, sox10* in turquoise module, commd5 in salmon, *tbx16l, pappa2* in brown and *tfr1a, aff2* in green; yet we also observed genes with opposite behaviour including *klf6a, irf2bp2a* in turquoise module, *sema4ab* in brown and *serinc2* in green module. No significant changes were observed between 48 and 72 hpf (data not shown), suggesting that both gene expression and chromatin accessibility were more stable by heart chamber formation.

**Figure 4.**
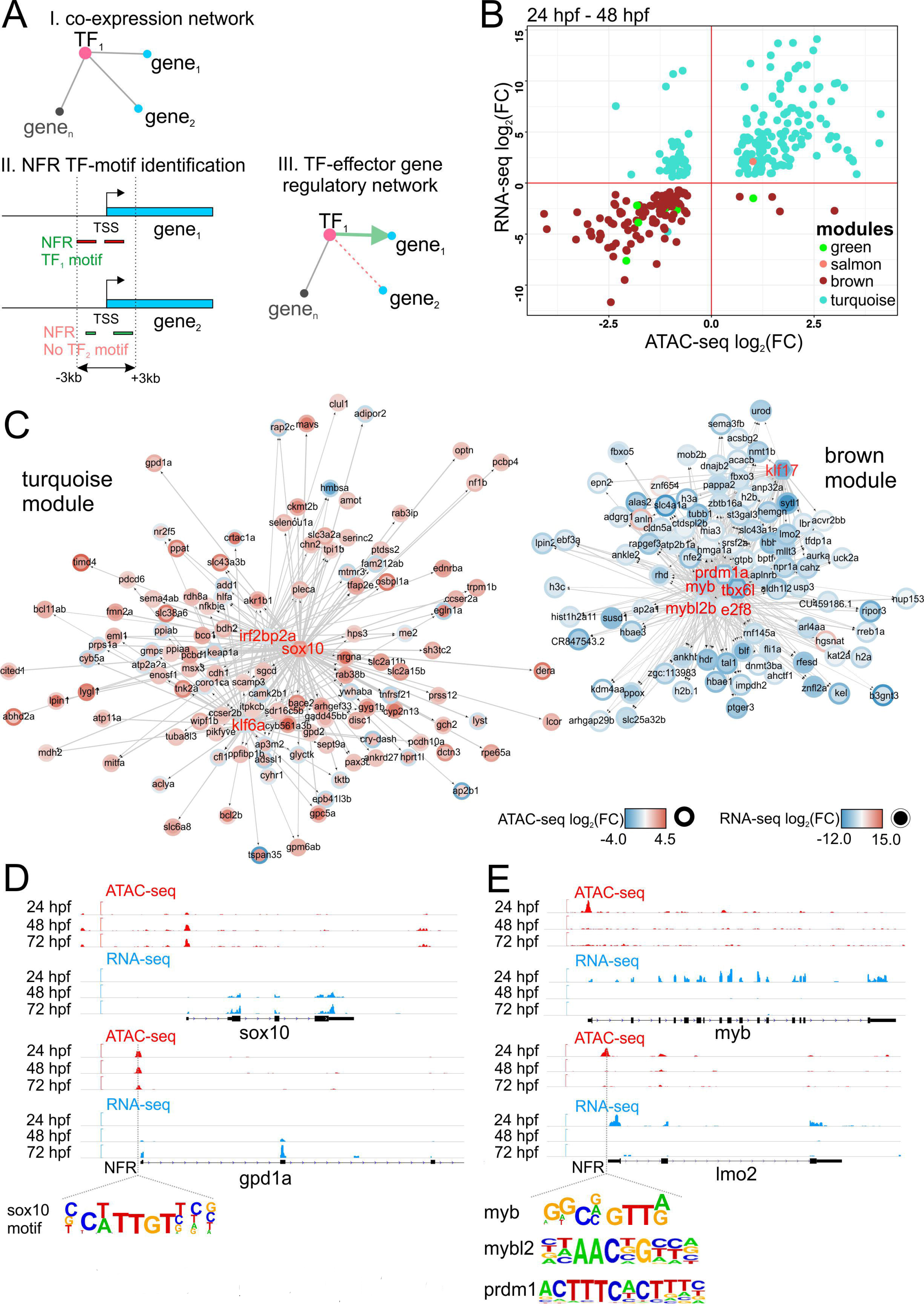
Dynamic regulatory networks of differentiating CMs. **(A)** Strategy used to establish gene-chromatin regulatory network. **(B)** Changes (log_2_FC) of gene expression compared to those in chromatin accessibility of cardiac module genes during heart development. Only significant (fdr < 0.05) genes are shown. **(C)** Regulatory networks of heart development. Arrows indicate the direction of interaction. Colours and the intensity of the circle edges indicate changes of chromatin accessibility, whereas those inside the circle show expression changes. Only significant (padj ≤ 0.05) genes are shown. Hub TFs are indicated in red font. **(D)** Visualization of ATAC-seq and RNA-seq read coverage of selected genomic regions related to turquoise module. **(E)** Visualization of ATAC-seq and RNA-seq read coverage of selected genomic regions related to brown module. Time points, NFRs and TF binding motifs within NFRs are indicated.

GO and pathway analysis (Croft et al. 2011) of turquoise regulatory network revealed that this module comprised genes involved in mitochondrial oxidation (*mdh2, gpd2*), carbohydrate metabolism (*rdh8a*) and ketone body metabolism (*bdh2*) (Fig. 4C-D, Supplement Table 6). We have identified *sox10, klf6a* and *irf2bp2a*, which were previously linked to zebrafish heart morphogenesis (Hill et al. 2017), as hub genes linked to their effector genes containing corresponding binding motifs in NFR localized in proximal promoter regions. As the vast majority of genes within the turquoise module exhibited significant increase in gene expression and chromatin accessibility within associated NFRs between 24 and 48 hpf, it suggests the presence of a metabolic switch that takes place in CM between those developmental stages. This agrees with previous reports showing that mitochondrial oxidative capacity and fatty acid oxidation potential increase along with CM maturation (Lopaschuk and Jaswal 2010).

Conversely, most of the genes assigned to brown module were downregulated from 48 hpf onwards along with the associated NFR chromatin accessibility (Fig. 4C). Pathway and GO analysis of brown module (Supplement Table 7) revealed the presence of genes implicated in embryonic haematopoiesis. Notably, we have identified a number of hub TFs including *myb* (*v-myb*) and *prdmla, mybl2, tbxl6l, e2f8, klf17* as well as their effector genes, such as *lmo2, tal1, alas2, slc4a1a* with profound roles in haematopoiesis (Fig. 4E) (Gering et al. 2003; Paw et al. 2003; Chan et al. 2009; Soza-Ried et al. 2010; Kotkamp et al. 2014). Moreover, ATAC-seq analyses revealed the enrichment of GATA, Fli1, ETS, ERG, and ETV motifs (Table 1) which belong to the regulatory network underlying the specification of hematopoietic and vascular lineages (Gottgens et al. 2002; Pimanda et al. 2007; Loughran et al. 2008; Kaneko et al. 2010). The brown module therefore represents the regulatory network leading to hematopoietic fate, whose suppression presumably promotes the development of CMs identity. Altogether, we identified regulatory networks leading to significant metabolic and cardiac/hematopoietic changes occurring in CMs during early heart morphogenesis (Supplement Table 8), which are regulated at both gene expression and chromatin levels.

### Disruption of cardiac TFs affects regulatory networks driving CM maturation

To further explore cardiac regulatory modules identified in our transcriptomic analyses and validate their importance in normal heart development, we utilized zebrafish mutants of cardiac TFs Gata5, Hand2 and Tbx5a, the disruption of which were previously linked to impaired migration of the cardiac primordia to the embryonic midline, reduced number of myocardial precursors and failure of heart looping, respectively (Reiter et al. 1999; Yelon et al. 2000; Garrity et al. 2002). RNA-seq and ATAC-seq were performed on CMs isolated from homozygous *gata5*^tm236a/tm236a^, *tbx5a*^m21/ m21^, *hand2*^s6,s6^ mutant 72 hpf embryos in Tg(*myl7*:EGFP) genetic background. Homozygous mutant embryos were selected based on their phenotypes of *cardia bifida* (*gata5*^tm236a/tm236a^, *hand2*^l6/s6^) or *heart-string* (*tbx5a*^m21/ m21^) (Fig. 5A).

**Figure 5.**
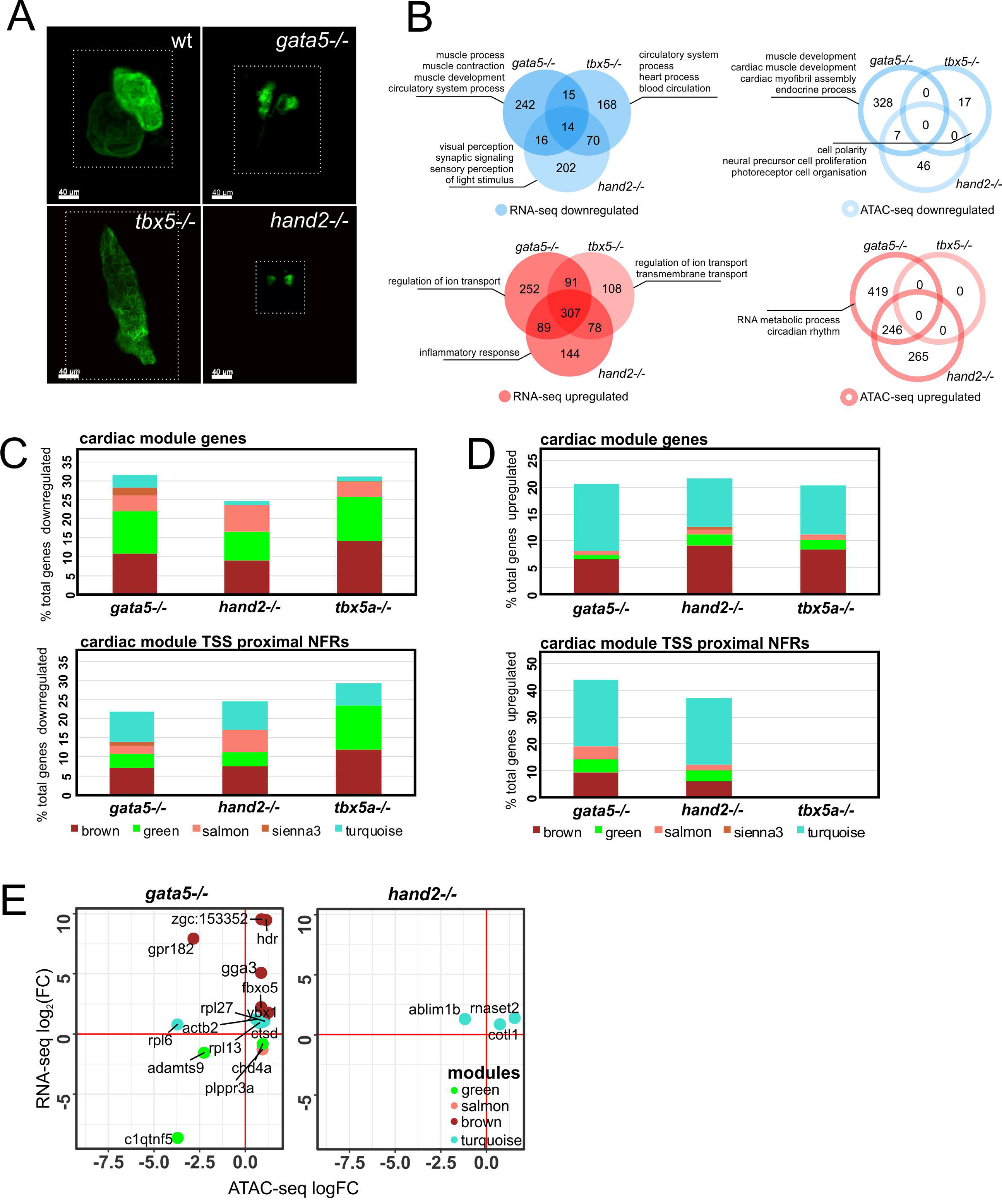
Loss-of-function mutations of cardiac TFs alters regulatory networks involved in heart development. **(A)** LSFM images of GFP-labeled CMs of wild-type and TF mutants zebrafish hearts at 72 hpf. Dotted line indicates exact area of the LSFM image. **(B)** Venn diagrams and GO enrichment analysis of TF-mutant downregulated (blue) and upregulated (red) genes and chromatin accessibility of proximal promoter NFRs (+/-3 kb of TSS), padj ≤ 0.05. **(C)** Percent distribution of cardiac module downregulated genes/proximal NFR chromatin accessibility as compared to total number of TF mutants downregulated genes/proximal NFR chromatin accessibility. **(D)** Percent distribution of cardiac module upregulated genes/proximal NFR chromatin accessibility as compared to total number of TF mutants upregulated genes/proximal NFR chromatin accessibility. **(E)** Cardiac module genes with differentially regulated expression and chromatin accessibility of proximal promoter NFRs (+/-3 kb of TSS) in *gata5, hand2*and *tbx5a* mutants.

RNA-seq analysis identified a number of genes which were differentially expressed (log_2_FC≠0, padj ≤ 0.05) in response to disruption of Gata5 (287 downregulated, 739 upregulated), Hand2 (288 downregulated, 618 upregulated) and Tbx5a (255 downregulated, 584 upregulated) (Fig. 5B, Supplement Table 9). Only a small overlap was observed between genes commonly downregulated in the three mutants (14 genes including *vcanb, bmp3* and *col18a1b*), whereas upregulated genes showed a larger overlap (307 genes e.g. *trim46, map4k6, mtf1*) between the three mutants. GO enrichment analysis of all TF-downregulated genes revealed the presence of biological processes related to muscle development, muscle function, heart process and sensory perception signalling; upregulated genes were enriched in biological processes related to ion transport and inflammatory response (Supplement Table 10).

On the other hand, changes within chromatin accessibility of NFRs localized in proximal promoter regions (+/- 3 kb of TSS) of mutants and wild-type embryos were generally less pronounced as compared to those at the gene expression level (Fig. 5B, Supplement Table 9). Moreover, loss of different TFs seems to have a variable effect on the chromatin structure, the largest of which seems to occur in *gata5*^tm236a/tm236a^ mutants (335 regions), where differentially represented proximal NFRs were associated with genes enriched in cardiac muscle development processes (Fig. 5B, Supplement Table 10). In *hand2^s6,s6^* mutants 53 regions were downregulated. Less pronounced chromatin changes could be identified in *tbx5a*^m21/ m21^ mutant (17 regions). Seven overlapping downregulated regions were identified between *gata5*^tm236a/tm236a^ and *hand2*^s6/s6^ mutants associated with *nkx1.21a, dmd, frzb, gpr4, vap*, whereas 246 overlapping upregulated regions were identified including those localized in the proximity of *nr4a1, mycbp2, irf2bp2a*, *rpl3*. No common differentially regulated proximal NFRs were, however, found across all three mutants.

In order to assess how the loss of function of critical cardiac TFs affects the regulatory networks of heart development, we further explored which fraction of mutant-downregulated genes contributes to the cardiac regulatory modules identified in wild-type analyses. We found that 31% (91 genes), 24% (71 genes) and 31% (79 genes) of total downregulated genes in *gata5*^m236a/tm236a^, *hand2*^s6,s6^, and *tbx5a*^m21/m21^ mutants were present in cardiac modules, mainly in the brown and green modules (Fig. 5C). Among the 14 genes which were commonly downregulated in all three mutants, we found 6 which belonged to cardiac modules, 4 of which belonged to green (*nid1b, papss2b, vcanb, bmp3*) and 2 to salmon (*plppr3a, spon1b*) modules. Genes including *vcan, plppr3a* and Bmp family were previously found to play a crucial role in heart morphogenesis and function (Marques and Yelon 2009; Kern et al. 2010; Chandra et al. 2018). Similar comparison performed for chromatin accessibility data revealed that 21% (73 regions), 24% (13 regions) and 29% (5 regions) of proximal NFRs which showed decreased accessibility in gata5^tm236a/tm236a^, *hand2*^s6/s6^, and *tbx5a*^m21/m21^ mutants were located within the proximal promoters of genes belonging to cardiac modules (Fig. 5C). We also explored mutant-upregulated genes and proximal NFRs and their contribution to cardiac modules (Fig. 5D). It showed that 20% (153 genes), 21% (134 genes) and 20% (119 genes) of total upregulated genes in *gata5*^m236a/tm236a^, *hand2*^s6/s6^, and *tbx5a*^m21/ m21^ mutants were present in cardiac modules, predominantly in the brown and turquoise modules. Consequently, the most prominent changes were observed for proximal NFRs in brown and turquoise modules, and 43 % (292 regions) and 37% (229 regions) of total upregulated NFRs contributed to cardiac modules in *gata5*^tm236a/tm236a^ and *hand2*^s6/s6^. No changes were observed in *tbx5a*^m21/m21^ mutants.

We further investigated the interactions between chromatin accessibility changes and gene regulation within cardiac modules in the three cardiac TF mutants. Hierarchical clustering revealed that the vast majority of either downregulated or upregulated cardiac module genes did not exhibit a similar regulation of NFR chromatin accessibility within their promoter regulatory regions (Supplemental Fig. 4). We observed that decrease in proximal promoter NFR were not correlated with gene expression downregulation, except for *c1qtnf5* and *adamts9*, the latter being a vcan-degrading protease required for correct heart development and cardiac allostasis (Kern et al. 2010) (Supplement Fig. 5 A, B). Similarly, only 10 genes including *hdr, gga3, fbxo5, rpl27, ybx1, actb2, cotl1, rnaset2* showed increase both in gene expression and NFR chromatin accessibility (Fig. 5E). Altogether, only 15 genes showed changes both in expression level and chromatin accessibility (either increasing or decreasing) in *gata5* mutant and 3 genes in *hand2* mutant, whereas no such genes were found in *tbx5a* mutant.

Taken together, we have identified a group of genes which were responsive to loss of Gata5, Hand2 and Tbx5a functions, among which, approximately one third belonged to cardiac regulatory networks. This suggests their crucial role in heart development and CM maturation downstream of these cardiac TFs. At the same time, it also provides a strong validation of the cardiac modules as gene regulatory networks underlying specific processes of heart development.

### Evolutionary conserved enhancers ensure proper heart development

One important observation was that gene expression changes in all three mutants were, to a large extent, uncorrelated with changes in chromatin accessibility, at least in proximal promoter regulatory regions. This led us to question whether loss of Gata5, Hand2, and/or Tbx5a cardiac TFs may cause global chromatin changes at genomic sites other than proximal gene promoters, and whether the observed changes in gene expression could be attributed to distal regulatory elements such as enhancers. To this end, we have identified distal NFRs (more than +/- 3 kb of TSS) and their differential accessibility between wild-type at 72 hpf and the mutants. We identified 59, 14 and 33 downregulated and 551, 321 and 2 regions upregulated (padj ≤ 0.05) in *gata5*^tm236a/tm236a^, *hand2*^s6/s6^, and *tbx5a*^m21/ m21^ mutants, respectively (Fig. 6A). Amongst downregulated regions, 1 region was in common between *gata5*^tm236a/tm236a^ and *tbx5*^2l/ m21^ mutants (Fig. 6B). On the other hand, much stronger overlap was observed between *gata5*^236a,tm236a^ and *hand2*^s6/s6^ mutants for upregulated regions (183 regions) whereas no overlap was found between *gata5*^m236a/tm236a^ and *tbx5*^m21/m21^. One region at chromosome 21 (Chr21:15013048-15013154) was commonly upregulated in all 3 mutants. To further explore the genomic localisation of differentially regulated distal NFRs and identify evolutionary conserved putative enhancers, we visualized them onto zebrafish genome and compared them with database of highly conserved non-coding elements (HCNE) between zebrafish and human (Engstrom et al. 2008) (Fig. 6C, Supplement Table 11). A total of 22 regions revealed conservancy between zebrafish and human genomic sequences among which 3 were downregulated in tbx5a and hand2 mutants, whereas 19 of them showed significantly increased accessibility in hand2 and gata5 mutants. Among 3 most downregulated HCNE were those localized on chromosome 1, between *hand2* and *fbxo8* genes (Chr1:37584384-37584724) as well as those localized in the introns of *ppp3ccb* (Chr10:20246264-20246845) and *akt7a* (Chr20:4714760-4715050) genes (Fig. 6D). We also identified HCNE-NFRs which increased in accessibility in *gata5* mutant (Chr1:8598642-8598893) and genomic region at chromosome 10 (Chr10:8580509-8581153) which was commonly regulated in *hand2* and *gata5* mutants (Fig. 6E). Therefore, we have determined a number of distal NFRs which accessibility is affected by mutations of cardiac TFs among which we pinpointed highly conserved NFRs serving as potential enhancers that may play key roles in heart development.

**Figure 6.**
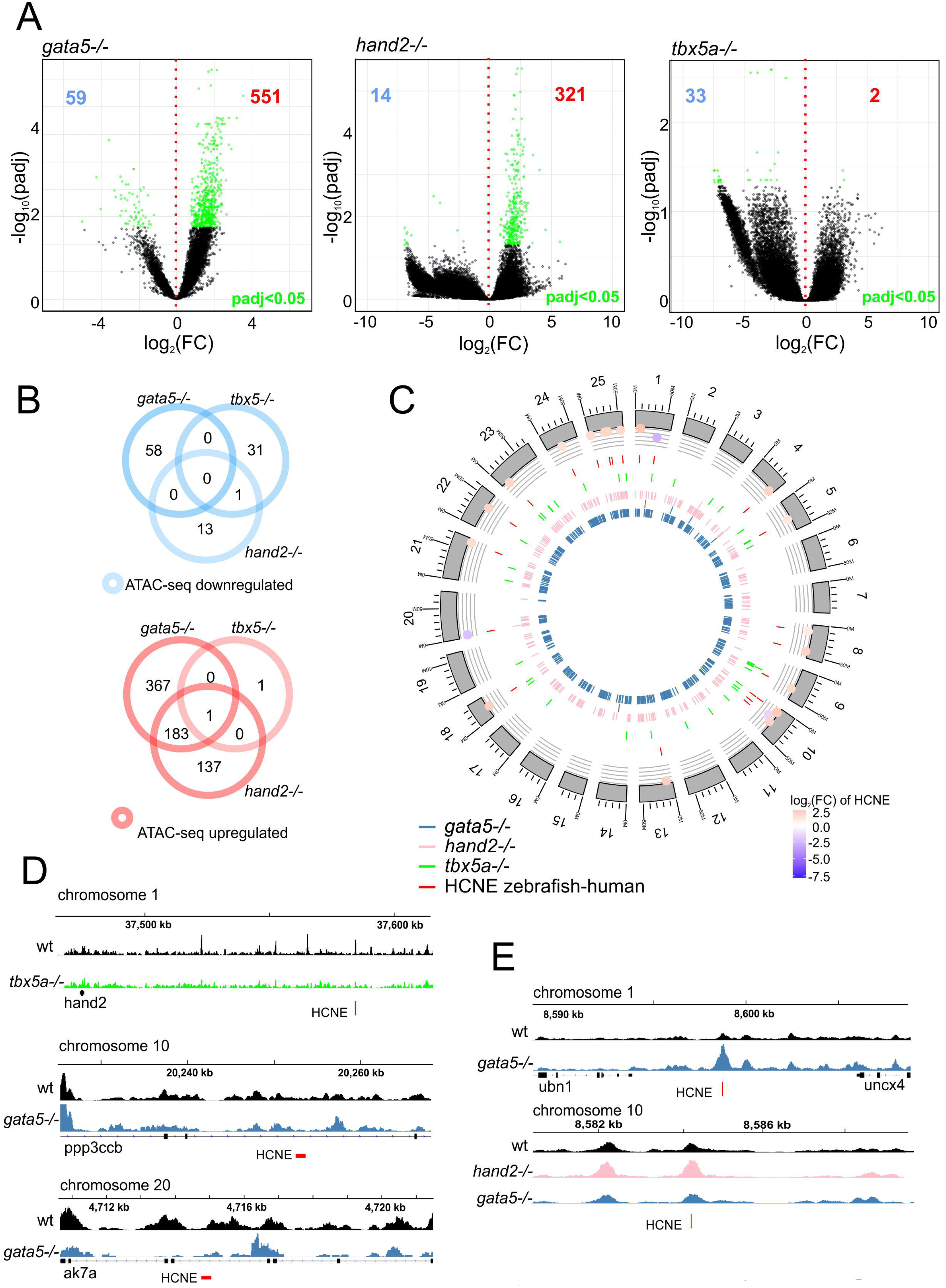
Identification of putative cardiac enhancers. **(A)** volcano plot of differentially accessible distal NFRs between wild-type and TF mutants at 72 hpf. padj ≤ 0.05 are indicated in green, number of downregulated NFRs is indicated in blue and upregulated in red. **(B)** Venn diagram of mutant down- and upregulated distal NFRs (more than +/- 3 kb of TSS), padj ≤ 0.05. **(C)** Graphical representation of differentially accessible distal NFRs genomic localization onto zebrafish chromosomes. NFRs overlapping with HCNE (+/- 500 bp) and their accessibility log2FC in comparison to wild-type is indicated, padj <0.05. **(D)** Genome track of ATAC-seq peaks for wild-type (black), *tbx5a-/-* (green) and *gata5-/-* (blue) for 3 most downregulated NFRs overlapping with HCNE (+/- 500 bp); **(e)** Genome track of ATAC-seq peaks for wild-type (black), *hand2-/-* (pink) and *gata5-/-* (blue) of 3 most upregulated NFRs overlapping with HCNE (+/- 500 bp).

## DISCUSSION

Heart development is a complex process involving multiple layers of interactions at molecular, cellular and tissue levels, with the former being controlled by a wide range of regulatory proteins including TFs, signalling proteins as well as epigenetic factors, such as histone and DNA modifications, chromatin remodelling and transcriptional enhancers. We used FACS to obtain CM-enriched cell fractions from developing heart during crucial events of heart morphogenesis. GFP-positive cells were sorted from transgenic Tg(*nxk2.5*:EGFP), Tg(*myl7*:EGFP) zebrafish embryos. In zebrafish, at 6-9 somite stage (∼12-14 hpf), *nkx2.5* expression only partially overlaps the anterior lateral plate mesoderm (ALPM) in its medial part (Schoenebeck et al. 2007), whereas at 17 somite stage (∼17-18 hpf) the most posterior *nkx2.5*+ cells of the bilateral cardiac primordia do not express *myl7*, a marker of terminal myocardial differentiation, suggesting the presence of *nkx2.5*+ cells that do not contribute to the myocardium (Yelon et al. 1999). This is in line with other studies in zebrafish, pinpointing the presence of specific *nkx2.5*+ second heart field (SHF) progenitors that give rise to the fraction of ventricular myocardium and outflow tract (OFT) (Guner-Ataman et al. 2013). Nevertheless, it has been shown that at prim-5 stage (24-30 hpf), *nkx2.5* is expressed both in ventricular and atrial myocardium exactly overlapping the expression of myl7 (Yelon et al. 1999). We applied an integrative approach combining transcriptomics (RNA-seq) and genome-wide chromatin accessibility maps (ATAC-seq). This strategy revealed several key observations. Firstly, the most prominent gene expression changes occurred between linear heart tube formation (24 hpf) and chamber formation (48 hpf). This major shift in molecular profile likely reflects the continuous process of CM differentiation throughout which progenitors are migrating and differentiate into CMs once they are incorporated into the growing heart tube (Kelly et al. 2014). Importantly, the genes which belong to sienna3 and turquois modules showed significant increase in expression between the two developmental stages. In particular, sienna3 genes were enriched in the largest number of GO terms related to cardiac function and contained at least three TFs known for their crucial roles in specification of CMs and their function in heart contraction (Singh et al. 2005; Singh et al. 2010; Laforest and Nemer 2011; Zhang et al. 2014; Pawlak et al. 2018), which suggests the prominent role of this network in CM differentiation and heart tube formation during this developmental period.

Secondly, we observed that both gene expression profile and chromatin landscape changed most significantly between 24 hpf and 48 hpf, suggesting that the changes in gene expression profiles during this stage were likely regulated at the chromatin level. Besides validating the biological relevance of our ATAC-seq dataset, this observation suggests that active chromatin remodelling occurs throughout development, and that the regions with differential accessibility represent *cis* regulatory hubs driving the biological processes associated with differentiating CMs.

Thirdly, the identified modules of co-regulated genes represent sub-networks underlying specific biological processes associated with heart development. Further integration of these gene networks with ATAC-seq data allowed us to link TFs to their putative target genes, which was supported by the enrichment of DNA binding motif for specific TFs within NFRs in proximal promoters of the genes within each particular module. Collectively, our analyses of the regulatory networks and their representative expression patterns revealed increased expression of genes defining CM structure and function, whereas the expression and proximal promoter chromatin accessibility of hematopoietic genes were suppressed during CM differentiation. A particularly intriguing finding was that sorted GFP-positive cells also expressed hematopoietic determinants at the earliest stage observed (24 hpf). These were strongly grouped into a single expression module (brown) and strongly correlated between gene expression dynamics and chromatin accessibility in proximal promoters that decreased between 24 hpf and 48 hpf. One possible explanation is that the expression of hemato-vascular genes was contributed by cells giving rise to the pharyngeal arch mesoderm which also express *nkx2.5* used as our selection marker. Nevertheless, our transcriptome profile, as well as microscopic observations, suggests that the majority of the GFP-positive cell populations are likely CMs, which is further supported by the finding that the highest-expressed genes were implicated in CM development and function. Another equally plausible hypothesis is that a group of cells exist within the pool of CM progenitors which possess alternative potential to become the blood or vascular lineage. Numerous evidences from mouse studies suggested the presence of bipotential cardiac progenitor populations which co-expressed cardiac and hematopoietic markers in the developing heart tube (Caprioli et al. 2011; Nakano et al. 2013; Zamir et al. 2017). The presence of hematopoietic markers in our experiment therefore suggests the presence of such cells in zebrafish and that, similar to mammals, the hematopoietic cell fate is suppressed with the progression of CM differentiation, a process which occurs between linear heart tube formation and chamber differentiation. To clearly distinguish between these possibilities, it would be necessary to obtain molecular profiles of individual cells so to determine whether hemato-vascular progenitors exist as a separate population expressing specific markers or rather, as a common progenitor population expressing both CM and hemato-vascular markers. Further, this also highlights the limitations of currently available marker genes, and calls for higher resolution analyses of gene expression in specific cell types which is possible with the single cell sequencing technology.

Finally, by performing parallel analyses in CMs isolated from mutants of cardiac TFs Gata5, Hand2 and Tbx5a, we uncovered changes in gene expression profiles and chromatin accessibility within cardiac regulatory networks. Comparing mutants and wild-type CMs, we observed only a minor correlation between changes in gene expression and chromatin accessibility within proximal promoter NFRs, suggesting that transcriptional regulation of genes involved in heart development might be affected by distal regulatory elements. Alternatively, changes in gene expression between wild-type and TF mutants could be related to impaired TF binding to constitutively accessible proximal NFRs. Moreover, due to the inability to distinguish mutant phenotype prior to 72 hpf, we could only perform mutant analyses at this developmental stage. This late stage of development means that we could not rule out the possibility that the effects we observe might be secondary in nature. Regardless that we could not provide definitive associations between distal regulatory elements and their target genes due to lack of chromatin interaction data, we identified a substantial number of gene-distal located NFRs which were altered in accessibility in mutants that may serve as potential distal transcriptional regulatory elements. Some of these elements were found to be highly conserved between zebrafish and human, suggesting that they might be critical developmental enhancers (Woolfe et al. 2005; Polychronopoulos et al. 2017).

Altogether, we characterized the dynamics of gene expression and chromatin landscape during heart development and identified genetic regulatory hubs driving biological processes in CMs at different stages of heart morphogenesis. We unravelled the alterations in the global transcriptional regulatory landscape resulting from disruptions to developmental program caused by the loss of cardiac TFs. Collectively, our study identified potential target genes and regulatory elements implicated in heart development and disease.

## METHODS

### Collection of embryos

Zebrafish transgenic lines Tg(*nxk2.5*:EGFP), Tg(*myl7*:EGFP) in AB wild-type and *gata5*^tm236a/+^ (Reiter et al. 1999), tbx5a^m21/+^ (Garrity et al. 2002), *hand2*^l6/+^ (Yelon et al. 2000) mutant background were maintained in the zebrafish facilities of the International Institute of Molecular and Cell Biology in Warsaw (License no. PL14656251), according to standard procedures and ethical practices recommended. Embryos were grown in embryo medium at 28°C, staged according to standard morphological criteria (Kimmel et al. 1995), and harvested at three different developmental stages: prim-5 (24 hpf), long-pec (48 hpf) and protruding-mouth (72 hpf).

### CM collection by fluorescence-activated cell sorting (FACS)

Cell suspension was prepared from 500 zebrafish embryos and larvae as previously described (Winata et al. 2013). Cells were verified microscopically for the viability by using trypan blue solution and used for further procedures when more than 90% of viable cell were obtained in the suspension. Fluorescent (GFP+) and non-fluorescent cells (GFP-) were sorted by using FACSAria II cytometer (BD Biosciences, USA). Cells were inspected for their relative size, granularity and relative fluorescence. Cell suspension obtained from wild-type embryo was used to assess the autofluorescence. GFP+ and GFP-fractions were verified for their viability by staining with propidium iodide (Sigma-Aldrich, USA) followed by FACS.

### qPCR

Total RNA was extracted from 100,000 GFP+ and GFP-cells obtained from zebrafish embryos by using TRIzol LS (Thermo Fisher Scientific, USA) according to the manufacturer protocol and followed by DNase I (Life Technologies, USA) treatment. Transcriptor first strand cDNA synthesis kit (Roche Life Science, Germany) was used to obtain cDNA. Relative mRNA expression was quantified by using FastStart SYBR green master mix on the Light Cycler 96 instrument (Roche Life Science, Germany) with specific sets of primers (Supplement Table 12).

### RNA-seq

For RNA sequencing 100,000 of GFP+ and GFP-cells from zebrafish embryos were sorted directly to TRIzol LS (Thermo Fisher Scientific, USA). After ethanol precipitation RNA was depleted of DNA by using DNase I treatment and purified on columns by using RNA Clean & Concentrator™-5 (Zymo Research, USA). RNA integrity was measured by RNA ScreenTape on the Agilent 2200 TapeStation system (Agilent Technologies, USA). RNA Integrity Number (RIN) was in the range from 8.5 to 10 for all the samples used for RNA-seq. Ribosomal RNA removal from 10 ng of total RNA was performed using RiboGone Kit (Clontech Laboratories, USA). cDNA synthesis for next-generation sequencing (NGS) was performed by SMARTer Universal Low Input RNA Kit (Clontech Laboratories, USA) as recommended by the manufacturer. DNA libraries were purified with Agencourt AMPure XP PCR purification beads (Beckman Coulter, USA) and DNA fragment distribution was assessed by using D1000 ScreenTape and Agilent 2200 TapeStation system (Agilent Technologies, USA). KAPA library quantification kit (Kapa Biosystems, USA) was used for qPCR-based quantification of the libraries obtained. Paired-end sequencing (2×75bp reads) was performed with NextSeq 500 sequencing system (Illumina, USA). The sequencing coverage was at least 75 million reads and 35 million reads for GFP+ and GFP-, respectively. GFP+ samples obtained from embryos at 24, 48 and 72 hpf were duplicated. Pearson correlation of biological replicates and read distribution over the zebrafish genome features were performed (Supplement Fig. 6 A, B)

### Assay for transposase-accessible chromatin with high throughput sequencing (ATAC-seq)

For ATAC-seq 60,000 of GFP+ cells from zebrafish embryos were sorted to Hank’s solution (1× HBSS, 2mg/mL BSA, 10 mM Hepes pH 8.0), centrifuged for 5 minutes at 500 × g and prepared for chromatin tagmentation as previously described (PMID: 24097267). NEBNext High-Fidelity 2 × PCR Master Mix (New England Biolabs, USA) and custom HP LC-purified primers containing Illumina-compatible indexes were used to prepare DNA sequencing libraries as previously described (Buenrostro et al. 2015). DNA libraries were purified with Agencourt AMPure XP PCR purification beads (Beckman Coulter, USA) and DNA fragment distribution was assessed by using D1000 ScreenTape and Agilent 2200 TapeStation system (Agilent Technologies, USA). KAPA library quantification kit (Kapa Biosystems, USA) was used for qPCR-based quantification of the libraries obtained. Paired-end sequencing (2×75bp reads) was performed with NextSeq500 sequencing system (Illumina, USA). The sequencing coverage was at least 90 million reads.

### Light sheet fluorescence microscopy (LSFM)

Embryos collected from transgenic lines Tg(*nxk2.5*:GFP) and Tg(*myl7*:GFP) were maintained in embryo medium containing 0.003% 1-phenyl-2-thiourea (PTU) to inhibit the development of pigment cells. Embryos collected from wild-type at 24, 48 and 72 hpf were mounted in 1% low-melting agarose (Sigma-Aldrich, USA) in a glass capillary. LSFM was used to perform optical sectioning of the cardiomyocytes containing GFP reporter. Images were analysed with Imaris 8 software (Bitplane, Switzerland).

### Bioinformatics analysis

Raw RNA-seq and ATAC-seq reads were quality checked using FastQC (0.11.5) (http://www.bioinformatics.babraham.ac.uk/projects/fastqc/) and MultiQC (1.1) (Ewels et al. 2016). Illumina adapters were removed using Trimmomatic (0.36) (Bolger et al. 2014). Reads matching ribosomal RNA were removed using rRNAdust (Hasegawa et al. 2014). Reads quality filtering was performed using SAMtools (1.4) (Li et al. 2009) with parameters -b -h -f 3 -F 3340 -q 30. RNA-seq reads were aligned to the zebrafish reference genome (GRCz10) using STAR (2.5) (Dobin et al. 2013) (Supplement Fig. 7). Bowtie2 (2.2.9) (Langmead and Salzberg 2012) was used to map ATAC-seq reads to the entire GRCz10 genome except *hand2*^s6/s6^ in which ∼200kb region spanning hand2 gene was excluded from the analysis due to large deletion carried by those mutants as previously described (Yelon et al. 2000) (Supplement Fig. 8). Read distribution was assessed with Picard (2.10.3). NFR regions were identified as previously described (Buenrostro et al. 2013). Peaks of chromatin open regions were called using MACS2 (2.1.0) (Zhang et al. 2008) with parameters --nomodel --shift -100 --extsize 200 --broad -g 1.21e9 -q 0.05 -B --keep-dup all. Enriched motifs in NFRs were identified using the HOMER findMotifsGenome tool with parameters findMotifs.pl modules/$modir/target.fa fasta modules/$modir -mset vertebrates -p 8 -S 200 -fastaBg modules/$modir/background.fa to check against vertebrates motif collection (Heinz et al. 2010). The background collection of sequences was constructed for each investigated gene module by taking complementing set of NFRs around TSSs of that module. Downstream bioinformatics analysis were performed in R 3.4 using following Bioconductor and CRAN (Huber et al. 2015) packages: GenomicFeatures (Lawrence et al. 2013), GenomicAlignments (Lawrence et al. 2013), DESeq2 (Love et al. 2014), pheatmap, LSD, ComplexHeatmap, biomaRt (Durinck et al. 2009), dplyr, WGCNA (Langfelder and Horvath 2008), ggplot2, reshape2, org.Dr.eg.db, clusterProfiler (Yu et al. 2012), ATACseqQC (Ou et al. 2018), ChIPseeker (Yu et al. 2015), DiffBind (Ross-Innes et al. 2012), ggbio (Yin et al. 2012). RNA-seq gene counts and ATAC-seq NFR read counts for all samples were transformed to regularized log (rld) (Supplement Table 13, 14). Gene network visualisation and statistical analysis of gene networks was performed using Cytoscape (Cline et al. 2007). Metascape was used to visualise the output of GO enrichment analysis (Tripathi et al. 2015).

## Supporting information

## DATA ACESS

RNA-seq and ATAC-seq data have been submitted to the NCBI Gene Expression Omnibus database (https://www.ncbi.nlm.nih.gov/geo/) under accession number GSE120238.

## ACKNOWLEDGMENTS

We thank D. Garrity for sharing *tbx5a*^m21/+^ and D. Yelon for *hand2*^s6,+^ zebrafish mutant lines and D. Stainier for sharing Tg(*nxk2.5*:GFP) and Tg(*myl7*:GFP). We thank V. Korzh and members of the Winata lab for fruitful discussions. This work was supported by EU/FP7 - Research Potential FISHMED, grant number: 316125, National Science Centre, Poland, OPUS grant number: 2014/13/B/NZ2/03863. MP is supported by Foundation for Polish Science and Ministry of Science and Higher Education, Poland and SONATA grant number 2014/15/D/NZ5/03421. MM is a recipient of the Postgraduate School of Molecular Medicine doctoral fellowship for the program “Next generation sequencing technologies in biomedicine and personalized medicine”.

## CONTRIBUTIONS

MP, KK, MM, JR performed bioinformatics and statistical analysis. MP, KK, AM collected embryos, performed *in vivo* experiments and collected biological material. MP and KAN prepared NGS libraries and performed RNA-seq and ATAC-seq. MP performed LSFM. LB and KP performed FACS. MP, CW,JR, KH contributed to genomic data analysis. MP, KK, MM, JR, PC, CW contributed to the design of the study and interpreted data. MP prepared the figures. MP and CW conceived the study and wrote the manuscript. CW is the corresponding senior author.

