## Supplementary material for "Dynamics of Cardiomyocyte Transcriptome and Chromatin Landscape Demarcates Key Events of Heart Development"

**24 hpf**

Genomic Annotation Summary:

| Feature | Frequency |
| --- | --- |
| Promoter (1-2kb) | 3.734 |
| Promoter (<=1kb) | 30.311 |
| Promoter (2-3kb) | 3.545 |
| 5' UTR | 0.269 |
| 3' UTR | 1.814 |
| 1st Exon | 0.002 |
| Other Exon | 9.496 |
| 1st Intron | 7.987 |
| Other Intron | 18.513 |
| Downstream (<=3kb) | 1.299 |
| Distal Intergenic | 23.026 |

**48 hpf**

Genomic Annotation Summary:

| Feature | Frequency |
| --- | --- |
| Promoter (1-2kb) | 3.595 |
| Promoter (<=1kb) | 33.850 |
| Promoter (2-3kb) | 3.342 |
| 5' UTR | 0.202 |
| 3' UTR | 1.274 |
| 1st Exon | 0.006 |
| Other Exon | 6.277 |
| 1st Intron | 8.680 |
| Other Intron | 19.300 |
| Downstream (<=3kb) | 1.128 |
| Distal Intergenic | 22.339 |

**72 hpf**

Genomic Annotation Summary:

| Feature | Frequency |
| --- | --- |
| Promoter (1-2kb) | 3.728 |
| Promoter (<=1kb) | 29.445 |
| Promoter (2-3kb) | 3.581 |
| 5' UTR | 0.168 |
| 3' UTR | 1.565 |
| 1st Exon | 0.002 |
| Other Exon | 7.221 |
| 1st Intron | 9.469 |
| Other Intron | 20.781 |
| Downstream (<=3kb) | 1.190 |
| Distal Intergenic | 22.845 |
