## Supplementary material for "Dynamics of Cardiomyocyte Transcriptome and Chromatin Landscape Demarcates Key Events of Heart Development"

| **Gene name** | **Accession number (RefSeq)** | **Primer name** | **Primer sequence** |
| --- | --- | --- | --- |
| *nkx2.5* | NM_131421 | nkx25_F1 | GCCATC CGGATC CTCTCT CT |
| *nkx2.5* | NM_131421 | nkx25_R1 | TTCCTG ACAAAA CCCGAT GTCTTT |
| *myl7* | NM_131329 | myl7_F2 | CCAGAG GAAACC ATCCTT GCT |
| *myl7* | NM_131329 | myl7_R2 | TGGTCA ACCTCT TCTGCT GTG |
| *neurog1* | NM_131041 | ngn1_F1 | CCAGCC CACCAA TAAGGT TATCAA |
| *neurog1* | NM_131041 | ngn1_R1 | TGGAGA CGCAGG TGGTTT TC |
| *gfp* | NA | gfp_R1 | GGCAAG CTGACC CTGAAG TT |
| *gfp* | NA | gfp_F1 | GGCGGA CTTGAA GAAGTC GT |

**Supplementary Table 4.** The list of oligonucleotides used for qPCR.
