## Supplementary material for "Dynamics of Cardiomyocyte Transcriptome and Chromatin Landscape Demarcates Key Events of Heart Development"

**SUPPLEMENT DATA**

**List of supplement figures**

Supplement Figure 1. Biological validation of FACS-sorted cell fractions.

Supplement Figure 2. Assessment of CM-specific and non-CM gene markers.

Supplement Figure 3. Cardiac module enriched HOMER-generated TF motifs.

Supplement Figure 4. Hierarchical clustering of TF-differentially regulated genes and TSS proximal NFRs.

Supplement Figure 5. Hierarchical clustering of commonly TF-regulated genes.

Supplement Figure 6. Quality control of RNA-seq.

Supplement Figure 7. Alignment statistics of RNA-seq from GFP+ and GFP- cells from zebrafish cardiomyocytes.

Supplement Figure 8. Alignment statistics of ATAC-seq from zebrafish cardiomyocytes at different stage of heart development.

**List of supplement tables**

Supplement Table 1. DESeq2 differentially expressed genes (DEG) GFP+ and GFP- cell fractions at 24, 48 and 72 hpf and DEG in GFP+ between developmental stages (padj $\leq$ 0.05). Numbers in columns are $log_2FC$ corresponding to compared conditions as indicated in table headers.

Supplement Table 2. Genomic annotation summary of ATAC-seq peaks at 24, 48 and 72 hpf.

Supplement Table 3. List of WGCNA modules and genes.

Supplement Table 4. GO enrichment analysis of WGCNA module genes. Padj value corresponds to p-value adjusted for multiple testing using Benjamini-Hochberg method.

Supplement Table 5. Differentially accessible gene-associated NFRs from proximal TSS regions (+/-3kb) at different stages of heart development. Numbers in columns are $log_2FC$ corresponding to compared conditions as indicated in table headers.

Supplement Table 6. DAVID GO enrichment analysis of turquoise module gene network.

Supplement Table 7. DAVID GO enrichment analysis of brown module gene network.

Supplement Table 8. Cytoscape-generated networks of turquoise and brown modules.

Supplement Table 9. DESeq2 DEG and differentially accessible proximal NFRs (+/- 3kb from TSS) between wild-type and TF mutant CMs (padj $\leq 0.05$). Numbers in columns are $\log_2$FC corresponding to compared conditions as indicated in table headers.

Supplement Table 10. GO enrichment analysis of DEG and differentially accessible proximal NFRs (+/- 3kb from TSS) between wild-type and TF mutant CMs. Padj value corresponds to p-value adjusted for multiple testing using Benjamini-Hochberg method.

Supplement Table 11. DESeq2 differentially accessible distal NFRs (more than +/- 3kb from TSS) between wild-type and TF mutants. Genomic coordinates are provided. HCNE_hu column indicates overlap (TRUE) with HCNE or lack of overlap (FALSE).

Supplement Table 12. qPCR primer list.

Supplement Table 13. DESeq2 regularized logarithm (rld) values for RNA-seq samples. Numbers in columns are rld corresponding to samples indicated in table headers.

Supplement Table 14. DESeq2 regularized logarithm (rld) values for ATAC-seq samples.

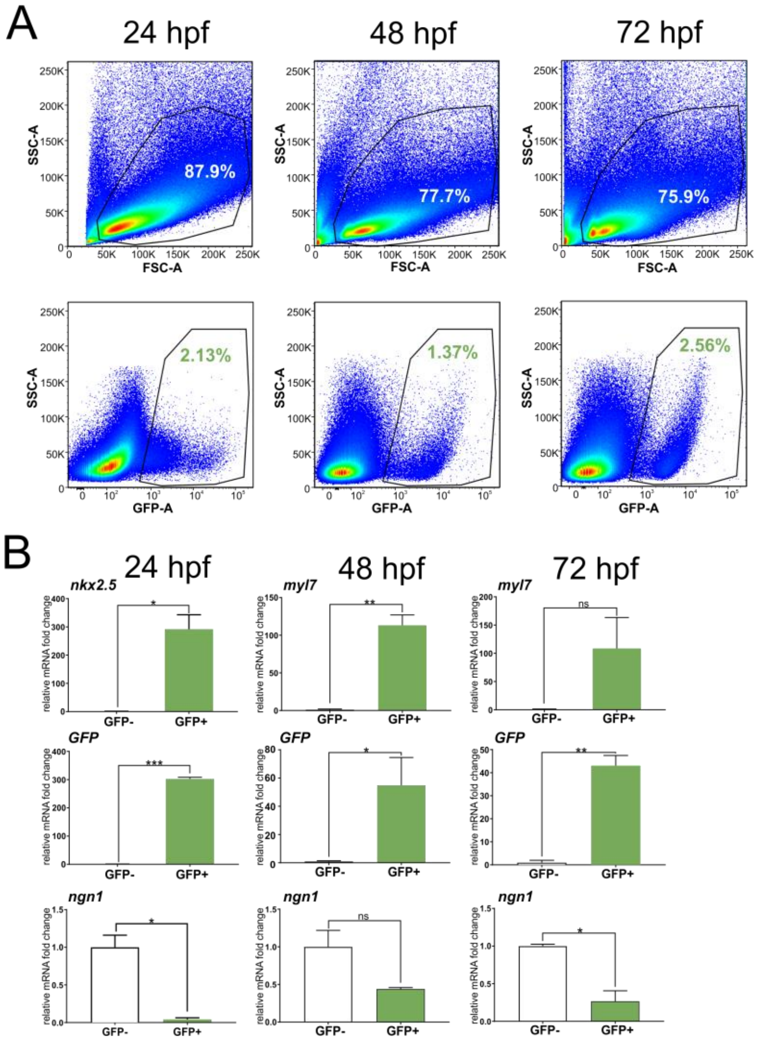


**Supplement Figure 1. Biological validation of FACS-sorted cell fractions. (A)** FACS analysis of cells obtained from developing zebrafish embryos. Forward scatter (FSC-A) vs side scatter (SSC-A) was used to find viable, single cell events; SSC-A vs GFP-A was applied to identify fluorescent cells. **(B)** Relative mRNA expression of nkx2.5, myl7, GFP and ngn1 in GFP+ and GFP- cells. **$p \leq 0.01$; ***$p \leq 0.005$; by unpaired t test. Data are presented as mean ± SD, n=2/group.

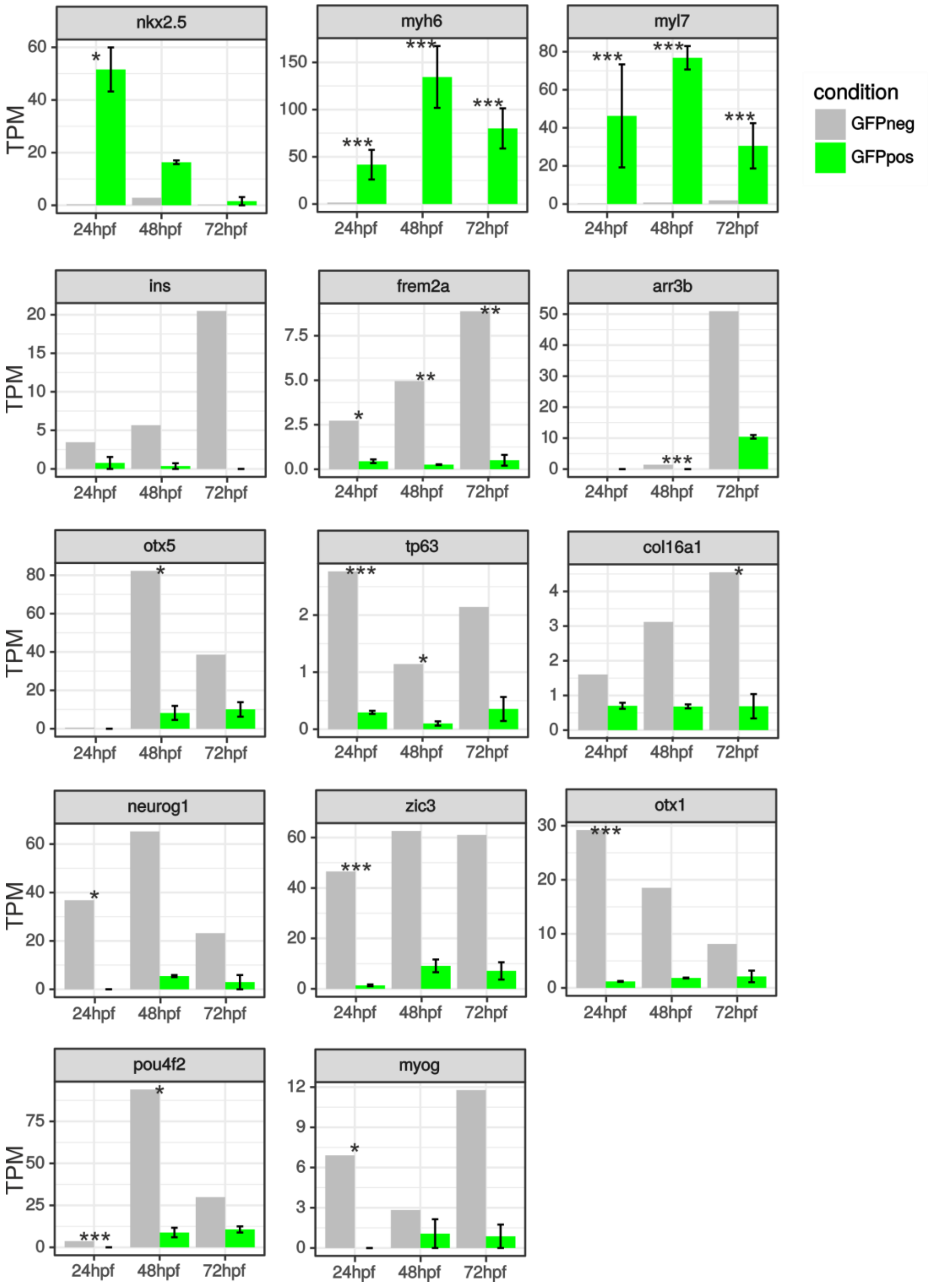


**Supplement Figure 2. Assessment of CM-specific and non-CM gene markers.** RNA-seq gene expression levels (transcripts per million, TPM) for CM-specific: nkx2.5 (early cardiac marker) and myl7, myh6, and non-CM markers: pancreas (ins), pharyngal arch (frem2a), retina (arr3b, otx5), skin (tp63, col16a1), neural system (neurog1, zic3, otx1), eye (pou4f2) and skeletal muscle (myog) are shown as mean ± SE, n=1/group for GFP- and n=2 for GFP+. **padj ≤ 0.01; *** padj ≤ 0.005; by Benjamini-Hochberg DESeq2 method.

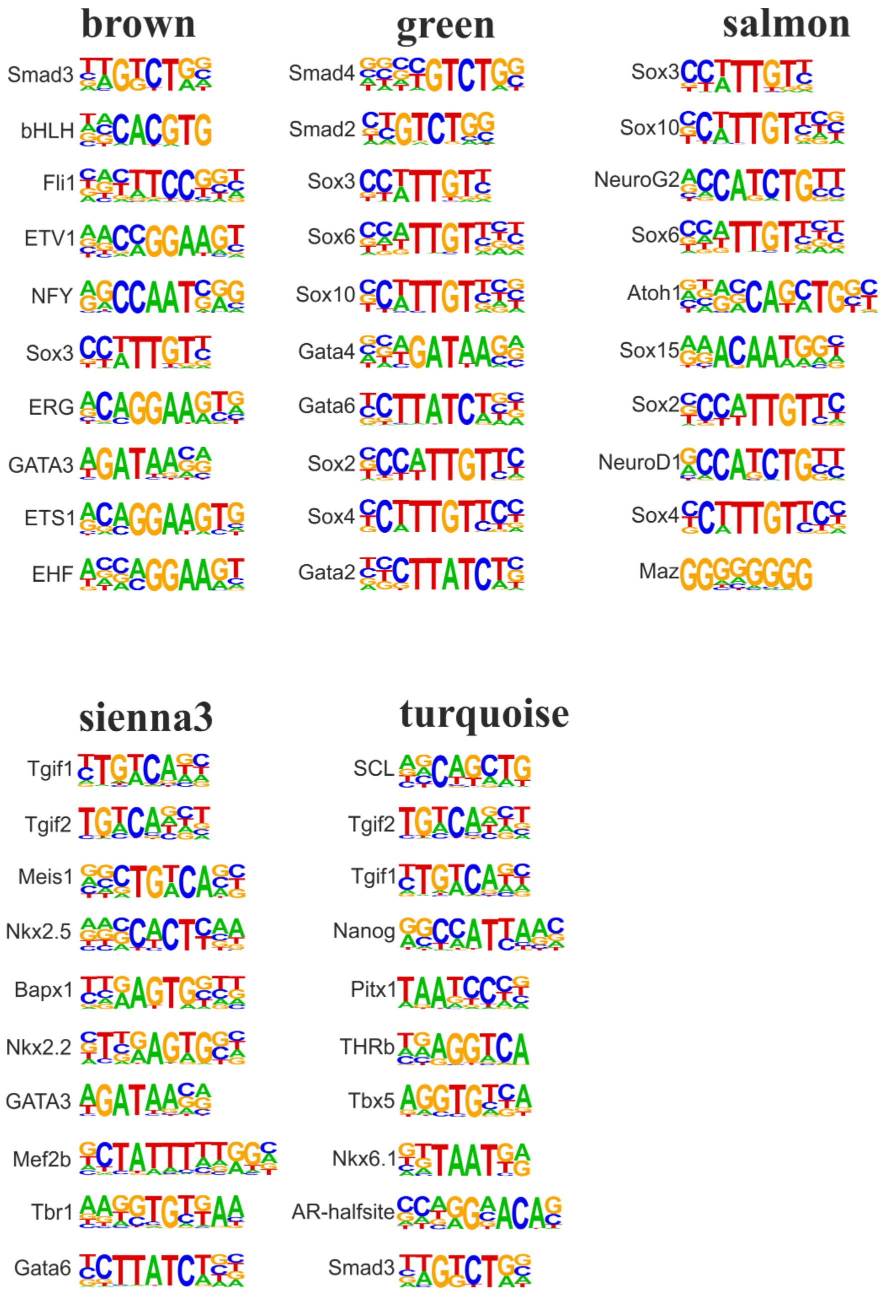


**Supplement Figure 3. Cardiac module enriched HOMER-generated TF motifs.** Graphical representation of Homer known motifs identified from cardiac modules. Ten motifs with p-value below 0.05 and highest number of target sequences are shown for each module.

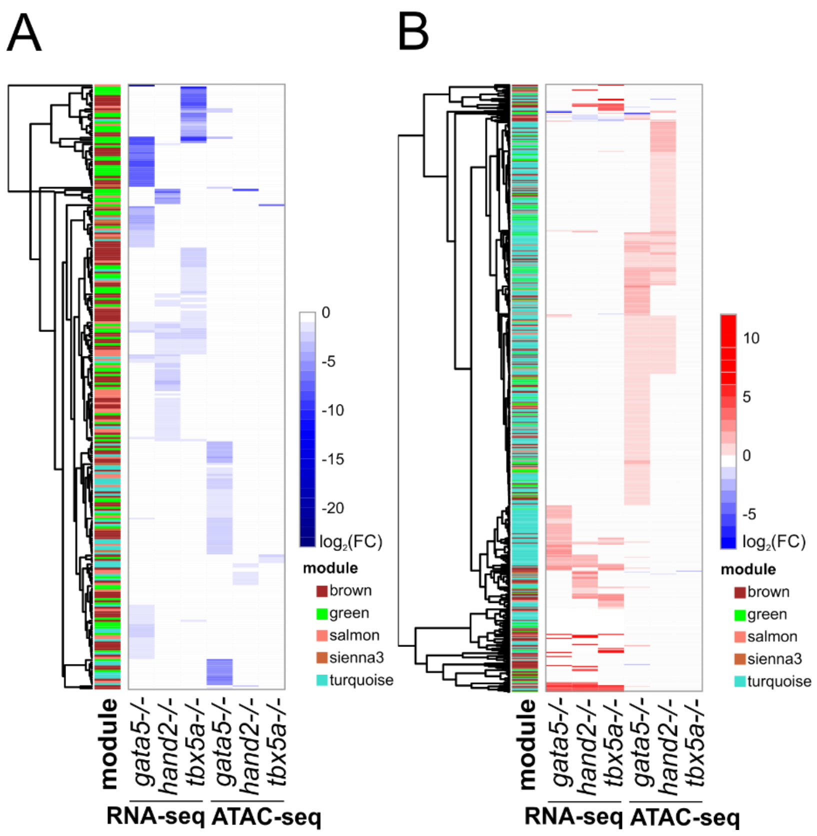


**Supplement Figure 4. Hierarchical clustering of TF-differentially regulated genes and TSS proximal NFRs. (A)** Hierarchical clustering of TF mutant downregulated genes and NFRs (+/- 3 kb of TSS) within cardiac regulatory modules. **(B)** Hierarchical clustering of TF mutant upregulated genes and NFRs (+/- 3 kb of TSS) within cardiac regulatory modules, padj $\leq 0.05$.

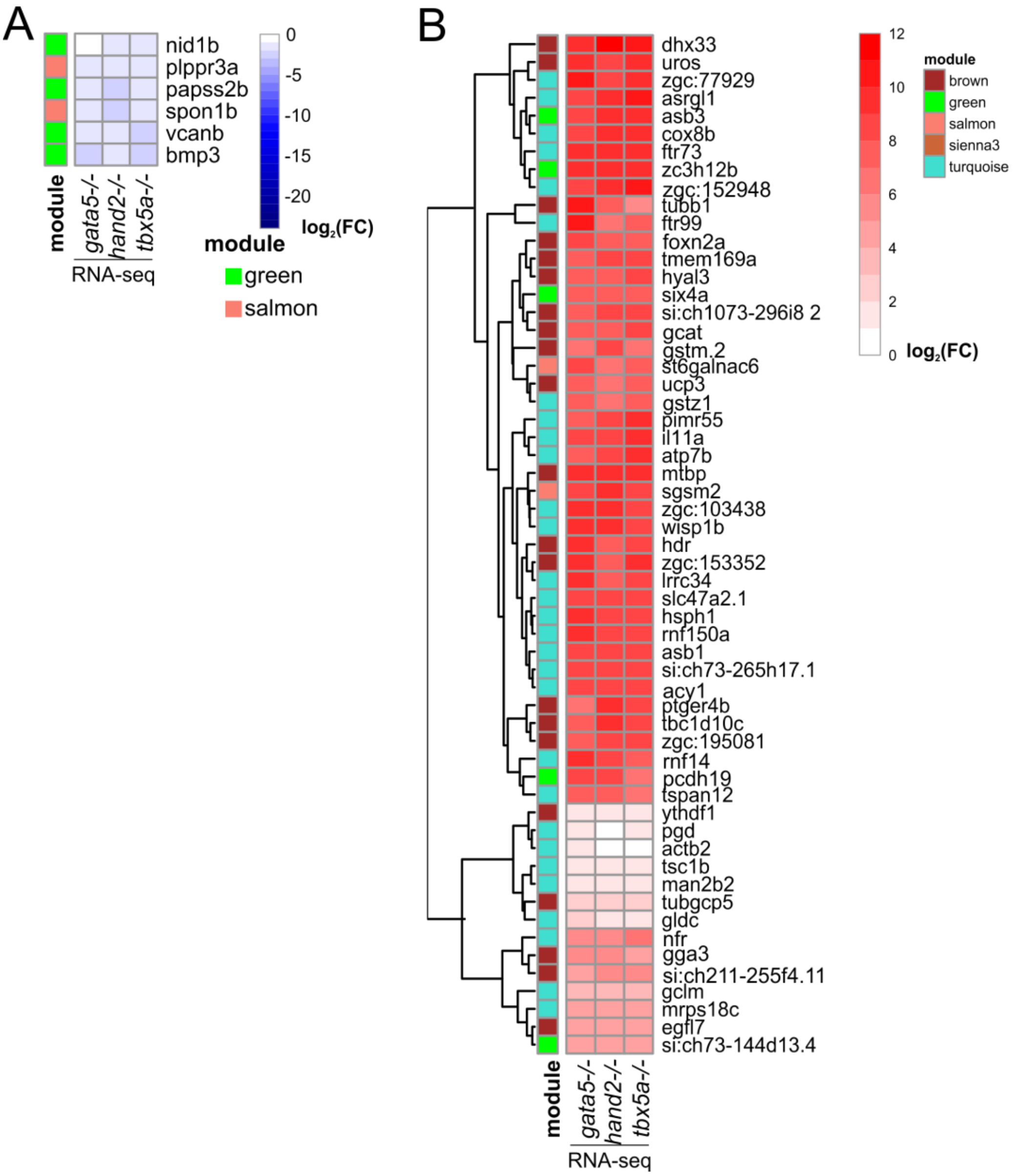


**Supplement Figure 5. Hierarchical clustering of commonly TF-regulated genes. (A)** Cardiac module genes commonly downregulated by TF mutants, padj ≤ 0.05. **(B)** Cardiac module genes commonly upregulated by TF mutants, padj ≤ 0.05.

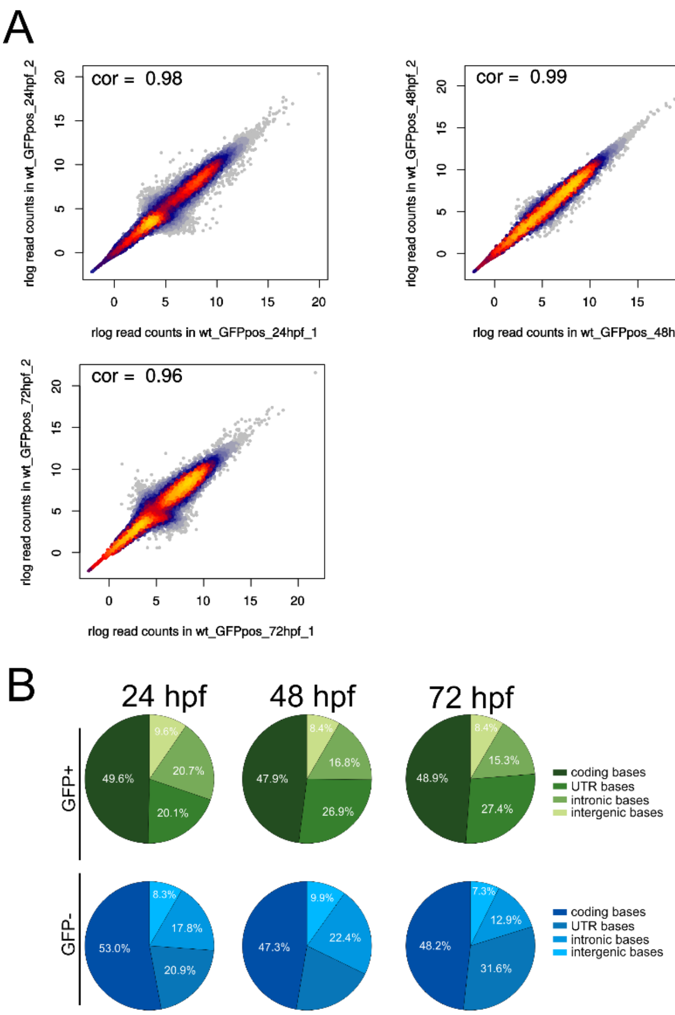


**Supplement Figure 6. Quality control of RNA-seq. (A)** Pearson correlation of normalized reads of RNA-seq biological replicates. **(B)** RNA-seq read distribution over zebrafish genome features.

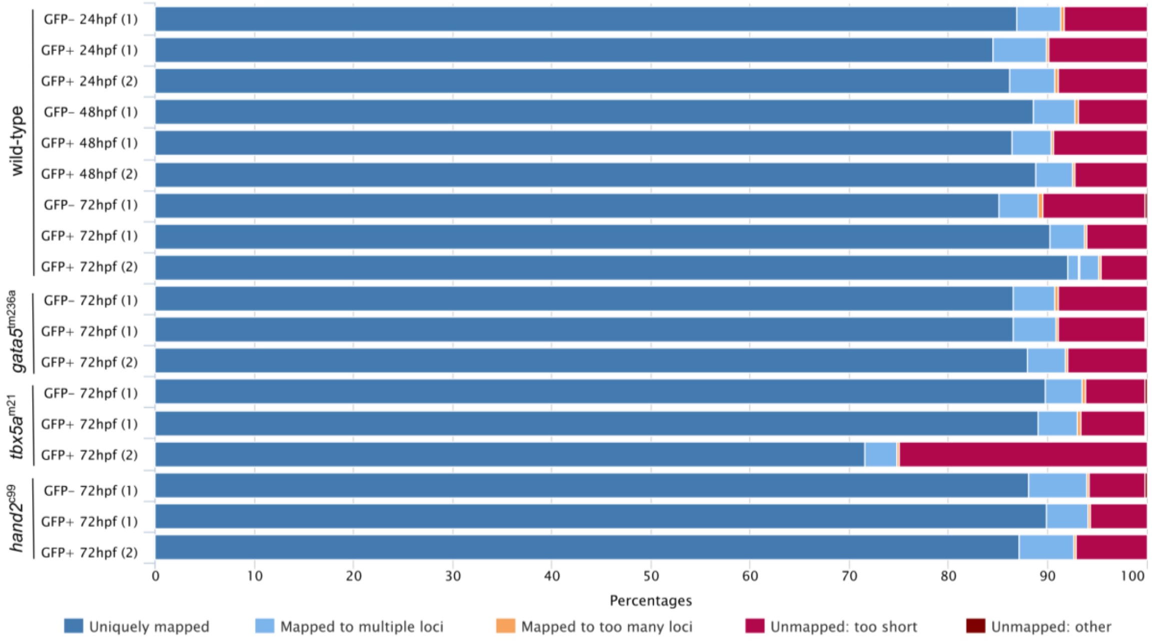


**Supplement Figure 7. Alignment statistics of RNA-seq from GFP+ and GFP- cells from zebrafish cardiomyocytes.**

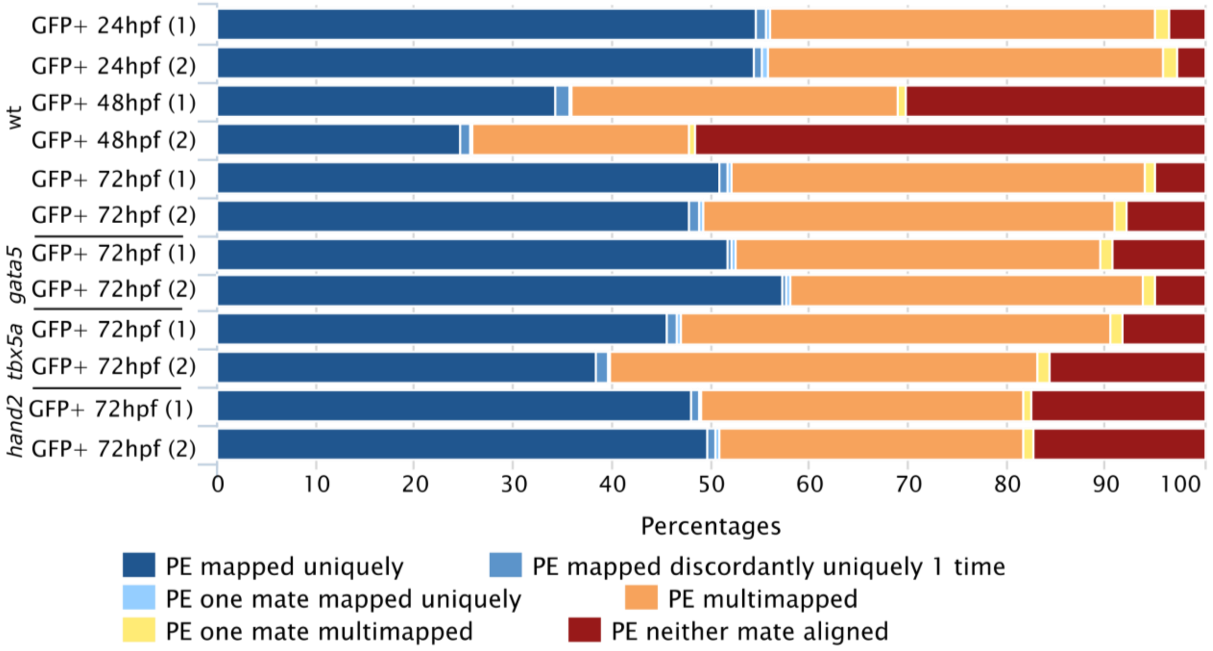


**Supplement Figure 8. Alignment statistics of ATAC-seq from zebrafish cardiomyocytes at different stage of heart development.** PE (paired-end).
